# New cell biological explanations for kinesin-linked axon degeneration

**DOI:** 10.1101/2021.12.23.473961

**Authors:** Yu-Ting Liew, André Voelzmann, Liliana M. Pinho-Correia, Thomas Murphy, Haydn Tortoishell, Jill Parkin, David M.D. Bailey, Matthias Landgraf, Andreas Prokop

**Affiliations:** The University of Manchester, Manchester Academic Health Science Centre, Faculty of Biology, Medicine and Health, School of Biology, Manchester, UK; Department of Zoology, University of Cambridge, Downing Street, Cambridge CB2 3EJ

**Keywords:** *Drosophila*, neurodegeneration, axons, microtubules, axonal transport, mitochondria, ROS

## Abstract

Axons are the slender, up to meter-long projections of neurons that form the biological cables wiring our bodies. Most of these delicate structures must survive for an organism’s lifetime, meaning up to a century in humans. Axon maintenance requires life-sustaining motor protein-driven transport distributing materials and organelles from the distant cell body. It seems logic that impairing this transport causes systemic deprivation linking to axon degeneration. But the key steps underlying these pathological processes are little understood. To investigate mechanisms triggered by motor protein aberrations, we studied more than 40 loss- and gain-of-function conditions of motor proteins, cargo linkers or further genes involved in related processes of cellular physiology. We used one standardised *Drosophila* primary neuron system and focussed on the organisation of axonal microtubule bundles as an easy to assess readout reflecting axon integrity. We found that bundle disintegration into curled microtubules is caused by the losses of Dynein heavy chain and the Kif1 and Kif5 homologues Unc-104 and Kinesin heavy chain (Khc). Using point mutations of Khc and functional loss of its linker proteins, we studied which of Khc’s sub-functions might link to microtubule curling. One cause was emergence of harmful reactive oxygen species through loss of Milton/Miro-mediated mitochondrial transport. In contrast, loss of the Kinesin light chain linker caused microtubule curling through an entirely different mechanism appearing to involve increased mechanical challenge to microtubule bundles through de-inhibition of Khc. The wider implications of our findings for the understanding of axon maintenance and pathology are discussed.

## Introduction

Axons are the long and slender processes of neurons which form the biological cables that wire the nervous system and are indispensable for its function. In humans, axons can be up to 2 metres long whilst displaying diameters of only 0.1-15μm (Prokop, 2020). Most of these delicate cellular processes must survive for an organism’s lifetime, meaning up to a century in humans. Unsurprisingly, mammals lose about 40% of their axon mass towards high age (Calkins, 2013; Coleman, 2005; Marner et al., 2003). This rate is drastically increased in hereditary forms of axonopathies (Prokop, 2021).

Of particular interest to this article are mutations of microtubule-binding motor proteins that cause axonopathies, of which the OMIM^®^ database (Online Mendelian Inheritance in Man^®^; Amberger et al., 2015) currently lists DYNACTIN 1 (ALS1, OMIM^®^ reference #105400), DYNEIN HEAVY CHAIN 1 (CMT2A1, #614228), KIF1B (CMT20, #118210), KIF5A (SPG10, #604187; ALS, #617921), KIF1A (SPG30, #610357; HSN2C, #614213), KIF1B (CMT2A1, #118210) and KIF1C (SPAX2, #611302); in the case of KIF1A, links to HSP and ataxias are likely to be added soon (Nicita et al., 2020). The listed motor proteins are actively involved in live-sustaining axonal transport of RNAs, proteins, lipids and organelles (Guedes-Dias and Holzbaur, 2019; Hirokawa et al., 2010). Genetic aberration of such transport is thought to lead to systemic collapse of axonal structure and physiology, hence axonopathy. However, we have little understanding of the concrete mechanisms leading to these pathologies.

To bridge this important knowledge gap, we took a novel approach based on two strategic decisions: Firstly, we used axonal microtubules (MTs) as our main readout. These MTs are arranged into loose bundles that run all along axons to form the essential highways for axonal transport and to provide a source of MTs for axon growth and branching at any life stage (Prokop, 2020). Accordingly, aberrations of MT bundles (presenting as gaps or areas of bundle disorganisation in the form of MT curling) are sensitive indicators of axonal pathology (Prokop, 2021). Mechanisms that help to maintain these MT bundles are starting to emerge (Hahn et al., 2019).

Our second strategic decision was the use of *Drosophila* primary neurons as a cost-effective and fast model system, where the complexity of mechanisms can be addressed with powerful genetics, in ways hard to achieve in vertebrate models (Prokop et al., 2013). For example, in this study alone, we used over 40 different mutations or transgenic constructs - some of them in genetic combinations to address functional redundancies or hierarchies (e.g. Beaven et al., 2015; Gonçalves-Pimentel et al., 2011; Koper et al., 2012). Loss-of-function analyses in *Drosophila* are facilitated by the fact that key factors, such as Kinesin heavy chain (Khc, kinesin-1), Kinesin light chain (Klc) or Milton, are each encoded by a single gene in *Drosophila*, as opposed to three, four or two in mammals, respectively. Furthermore, genetic and pharmacological tools are readily available to manipulate virtually any genes in question - and all these functional approaches can be combined with efficient and well-established readouts for axonal physiology and MTs (Hahn et al., 2016; Prokop et al., 2013; Prokop et al., 2012; Sánchez-Soriano et al., 2010).

Here we demonstrate that the losses of three motor proteins cause MT curling: the Kif5A/B homologue Kinesin heavy chain (Khc), the Kif1A homologue Unc-104, and Dynein heavy chain (Dhc/DYNC1H1). We find that all three are required for axonal transport of mitochondria and synaptic proteins. Focussing on Khc and employing available means to dissect its various sub-functions, we identified two mechanisms linking to MT curling: Firstly, loss of Khc/Milton/Miro-mediated transport causes harmful reactive oxygen species (ROS) likely linking to mitochondrial transport. Secondly, loss of the Kinesin light chain linker appears to cause de-inhibition of Khc as a condition that we find to cause MT curling. Both mechanisms align with the recently proposed ‘dependency cycle of local axon homeostasis’ as a conceptual model of axonopathy (Prokop, 2021).

## Results

### Loss of Khc, Unc-104 or Dhc cause axonal MT curling

To assess whether loss of motor protein function impacts on axonal MT organisation, we tested mutant alleles for axonal transport-related motor proteins: (a) Dynein heavy chain (Dhc) is an obligatory component of the dynein/Dynactin complex and essential for most, if not all, MT-based retrograde transport (Reck-Peterson et al., 2018); (b) Klp64D is an obligatory subunit of heterodimeric kinesin-2 (KIF3 homologue) reported to mediate anterograde axonal transport of actylcholine-related synaptic enzymes or olfactory receptors (Baqri et al., 2006; Jana et al., 2021; Kulkarni et al., 2017; Ray et al., 1999); (c) the PX-domain-containing type 3 kinesin Klp98A (KIF16B homologue) was shown to mediate autophagosome-lysosome dynamics and endosomal Wingless transport in non-neuronal cells but is also strongly expressed in the nervous system (Mauvezin et al., 2016; Witte et al., 2020; flybase.org: FBgn0004387); (d) the PH-domain-containing type 3 kinesin Unc-104 (Kif1A homologue) is essential for synaptic transport in *Drosophila* axons (Pack-Chung et al., 2007; Voelzmann et al., 2016); (e) Kinesin heavy chain/Khc is the sole kinesin-1 in *Drosophila* (Kif5A-C homologue) involved in multiple transport functions in *Drosophila* neurons (see details below; e.g. Bowman et al., 2000; Gindhart et al., 2003; Glater et al., 2006; Loiseau et al., 2010; Rosa-Ferreira et al., 2018; Saxton et al., 1991).

To assess potential roles of these motor proteins in MT regulation, we cultured primary neurons obtained from embryos lacking these gene functions (see Methods; Fig.1) and analysed them at 5DIV (days *in vitro*). MT curling phenotypes (where bundles deteriorate into curled, intertwined, crisscrossing MTs; curved arrows and enlarged insets in Fig.1B,F,G) occurred as a moderate phenotype in Dhc-deficient neurons and were prominent in neurons lacking Khc or Unc-104 (Fig.1H).

**Fig.1.**
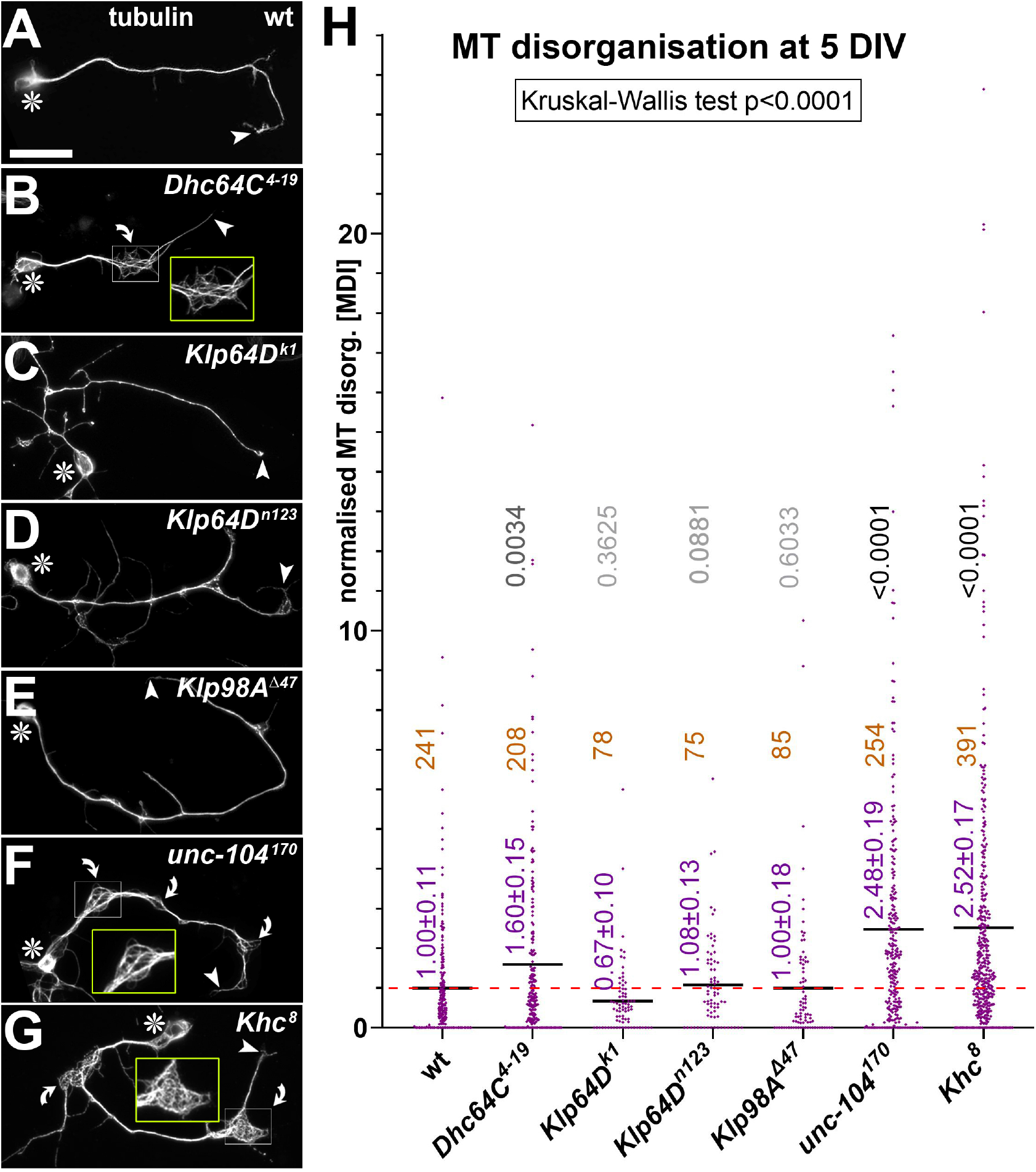
Deficiencies of three motor proteins cause MT curling. **A-G**) Examples of neurons of different genotype (indicated top right) and stained for tubulin at 5DIV; asterisks indicate cell bodies, arrow heads axon tips, curved arrows areas of MT curling, white rectangles shown as twofold magnified, yellow emboxed insets; scale bar in A represents 20μm in all images. **H**) Quantification of MT curling phenotypes measured as MT disorganisation index (MDI) and normalised to wild-type controls (red stippled line); mean ± SEM is indicated in blue, numbers of analysed neurons in orange, results of Mann Whitney rank sum tests are shown in grey/black.

To assess whether MT curling phenotypes were accompanied by transport defects, we analysed additional sub-cellular markers in *Khc^8^*, *unc-104^170^* and *Dhc64C^4-19^* homozygous mutant neurons. First, using the pre-synaptic protein Synaptotagmin (Syt) as an indicator of vesicular transport (Voelzmann et al., 2016), we found reduced presynaptic spots within axons of neurons mutant for any one of these three motor proteins, suggesting they all contribute to axonal vesicular transport (Fig.2A-D,O). Second, the axonal number and distribution of mitochondria (visualised with mitoTracker; Klionsky et al., 2012), is significantly reduced in neurons lacking either Khc, Unc-104 or Dhc64C function (Fig.2H-K,P). Third, in *Khc^8^* mutant neurons, we also assessed the distribution of endoplasmic reticulum (ER) using the genomically tagged *Rtnl1-GFP* allele. In wild-type neurons, ER is distributed evenly along the entirety of the axon (Fig.S1A); loss of Khc does not affect this distribution, but about three quarters of neurons show an abnormal accumulation of Rtnl-1::GFP*-* labelled ER material at their tips (Fig.S1B,C; details in legend).

**Fig.2.**
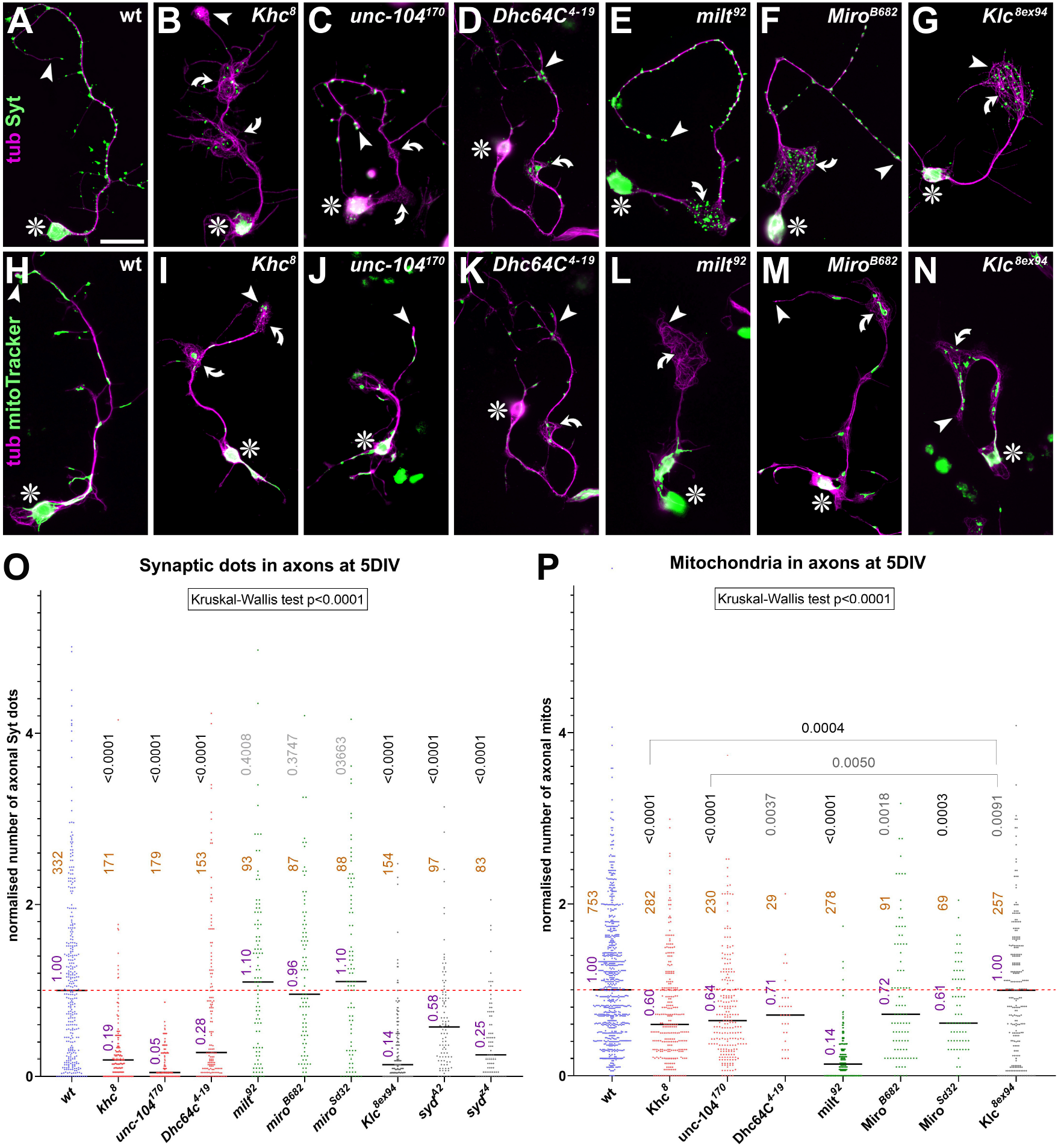
Impacts of motor protein and linker mutations on numbers of axonal mitochondria and synaptic spots. **A-N**) Examples of neurons of different genotype (indicated top right) and stained at 5DIV for tubulin (tub, magenta) and either Synaptotagmin (Syt, green in A-G) or with mitoTracker (green in H-N); scale bar in A represents 20μm in all images. **O,P**) Quantification of axonal numbers of Syt-positive spots (O) or mitochondria (P), all normalised to wild-type controls (red stippled line); medians are indicated in blue, numbers of analysed neurons in orange, results of Mann Whitney rank sum tests are shown in grey/black.

In conclusion, the three motor proteins that display MT curling are also required for normal axonal transport of synaptic vesicles and mitochondria in *Drosophila* primary neurons. In addition, at least Khc is also required for the normal axonal distribution of ER.

### Khc displays strong maternal effects

Of these three motor proteins, we performed more detailed analyses on Khc because many genetic tools are available for the systematic dissection of its various functions (Fig.3B). First, to validate its MT-related phenotypes, we tested additional mutant alleles (*Khc^27^* and *Khc^8^* in homozygosis or over deficiency) as well as RNAi mediated knockdown of Khc (via pan-neuronal *elav-Gal4*). In all cases we found that the MT curling phenotypes were equally present at 5DIV (Fig.S2A,B,D).

**Fig.3.**
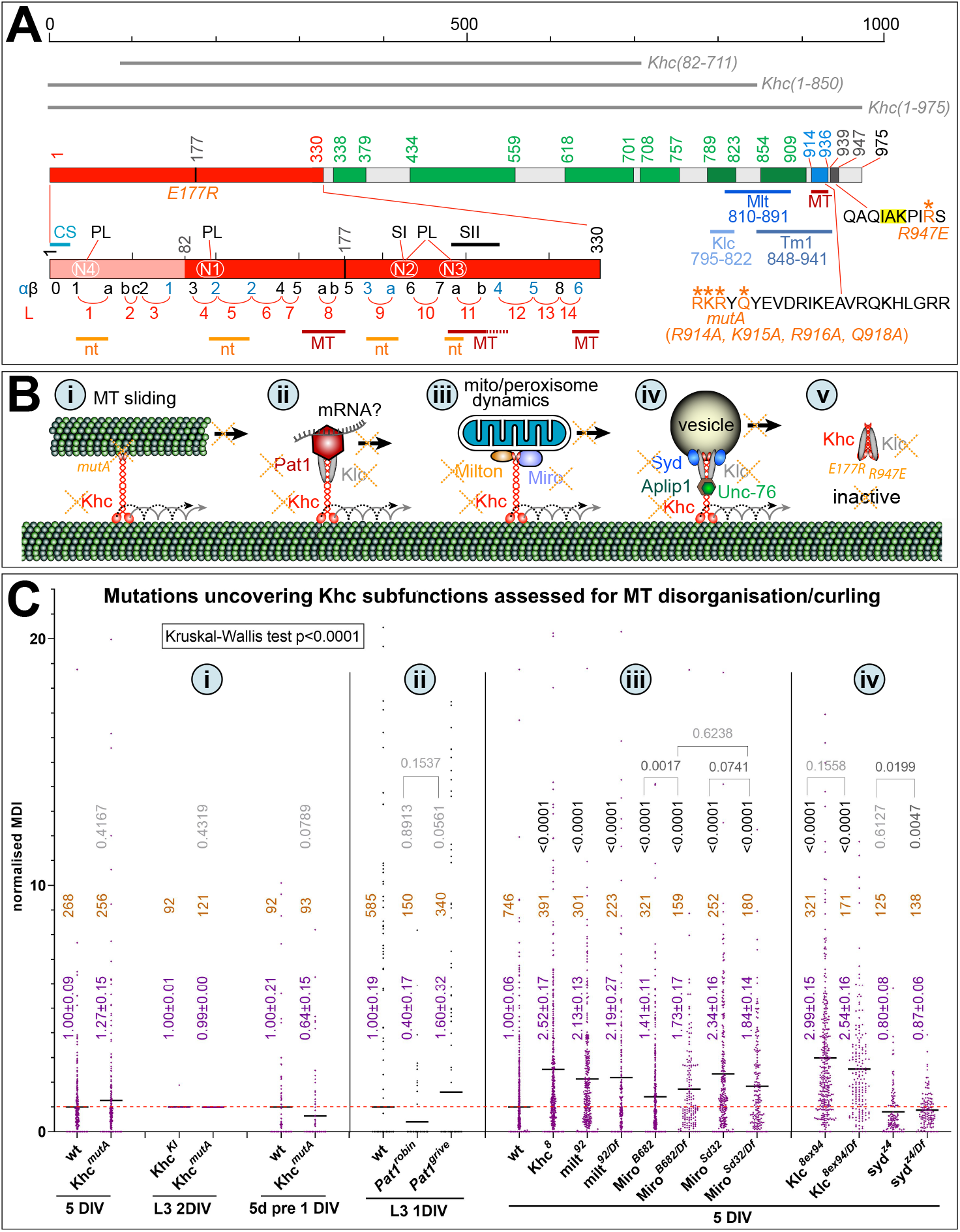
Assessing contributions of Khc subfunctions to MT regulation. **A**) Schematic representation of *Drosophila* Khc drawn to scale. Domains are colour-coded and start/end residues are indicated by numbers: motor domain (red; according to Sablin et al., 1996), coiled-coil domains required for homo- and/or heterodimerisation (green; as predicted by Ncoils in ensembl.org), the C-terminal ATP-independent MT-binding motif (blue; according to Winding et al., 2016), and the C-terminal auto-inactivation domain (dark grey; according to Kaan et al., 2011); grey lines above the protein scheme indicate the three expression constructs used in this study; below the protein scheme further details are shown: the sequence of the C-terminal MT-binding domain (*mutA* mutations indicated in orange; Winding et al., 2016), the sequence of the auto-inactivation domain (indicating the IAK motif and R947E mutation; Kelliher et al., 2018), the binding areas (darker green coiled-coils) of Klc (according to Veeranan-Karmegam et al., 2016), Mlt (known to overlap with Klc; Glater et al., 2006; Verhey et al., 1998) and Tropomyosin 1 (Dimitrova-Paternoga et al., 2021), and the two-fold enlarged motor domain. The secondary structure of the motor domain is indicated below (α helices in black, β sheets in blue, loops/L in red); this map was generated by matching the resolved structure of Khc (UniProt code: P17210, PDB id 2y65) with descriptions of the kinesin consensus (Sablin et al., 1996); regions/motifs that bind ADP/ATP (nt, orange; according to Cao et al., 2017; Gigant et al., 2013; Sablin et al., 1996) and/or MTs (dark red; according to Hunter and Allingham, 2020) are also indicated below; N1-4 in the motor domain indicate highly conserved motifs (according to Sablin et al., 1996); abbreviations above the motor domain indicate the locations of the cover strand (CS; according to Budaitis et al., 2021), P-loops (PL) and switch domains I and II (SI, SII; according to Cao et al., 2017; Gigant et al., 2013; Sablin et al., 1996). The N-terminal deletion of the above *Khc*(*82-711*) construct is shown in pink: it does not affect MT-binding sites, but it removes the cover strand (known to affect kinesin’s MT affinity and processivity; Budaitis et al., 2021) and the first P-loop (with potential impact on the ATP/ADP cycle); it might also affect the behaviour of the second P-loop which was shown to accelerate Khc movement when harbouring the T94S mutation (Cao et al., 2017; Higuchi et al., 2004). **B**) Schematic representation of some sub-functions of Khc (details and abbreviations in main text; red and stippled black lines indicate processive transport; for further sub-functions see Discussion): via a C-terminal MT-binding domain Khc can slide MTs (i), associating with Pat1 (and potentially Klc) it is expected to transport non-vesicular cargoes including mRNA (ii), with Milt and Miro organelle transport (iii), and with a protein complex containing Klc and Syd vesicular transport (iv); in the absence of such associations Khc is auto-inhibited and detaches from MTs assisted by Klc (v); to interfere with these subfunctions in this study, different genes were genetically removed (orange crosses) or specific *Khc* mutant alleles used (italic orange text). **C**) Quantified effects on MT curling caused by specific mutations affecting Khc sub-functions (numbers in grey circles indicate which function in A is affected): MT curling is quantified as MT disorganisation index (MDI) normalised to wild-type controls (red stippled line); bars at bottom indicate type of culture (‘5 DIV’, embryonic neurons 5 days *in vitro*; ‘L3 1/2 DIV’, late larval neurons 1/2 days *in vitro*; ‘5d pre 1 DIV’, embryonic neurons pre-cultured for 5 days and cultured for 1 day); mean ± SEM is indicated in blue, numbers of analysed neurons in orange, results of Mann Whitney rank sum tests are shown in grey/black.

However, the MT phenotypes were not evident at earlier stages in *Khc^8/Df^* mutant neurons, either at 6 hours *in vitro* (HIV) or at 3DIV (Fig.S2B,D), suggesting that phenotypes either accumulate gradually (as observed upon loss of Efa6; Qu et al., 2019) or are masked by perdurance of maternal product (meaning wild-type *Khc* gene product deposited in the egg by the heterozygous mothers; Prokop, 2013).

To distinguish between these two possibilities, we used a pre-culture technique where neurons are kept in centrifuge tubes for 5 days to deplete maternal gene product before plated in culture (Prokop et al., 2012; Sánchez-Soriano et al., 2010). Such pre-cultured neurons displayed prominent MT curling already at 1DIV (Fig.S2C,E), arguing that Khc has a prominent maternal contribution that persists for more than 3 days. Similar observations were made with the *Khc^1ts^* mutant allele (details in Fig.S2C).

### MT sliding functions of Khc do not link to MT curling

A C-terminal MT-binding site enables Khc to cross-link MTs and move them against each other (Fig.3Bi; Andrews et al., 1993; Jolly et al., 2010; Lu et al., 2013; Lu et al., 2015; Winding et al., 2016). We hypothesised that Khc might contribute to MT bundle maintenance by using its MT sliding function, for example by shifting MTs to achieve even distribution along axons. The Khc sliding function is selectively inhibited by the genomically engineered, lethal *Khc^mutA^* allele that abolishes C-terminal MT binding without interfering with other linkers or autoinhibition of Khc (Fig.3A; Winding et al., 2016).

To test whether Khc-mediated sliding contributes to MT bundle regulation, we cultured *Khc^mutA^* mutant neurons in different ways: embryo-derived neurons were cultured for 1DIV after 5d pre-culture or for 5DIV without pre-culture, and neurons from larval brains were cultured for 2DIV. In all cases, these neurons failed to show enhanced MT curling (Fig.3Ci), suggesting that the *Khc* mutant phenotype is not caused by the loss of its MT sliding function.

### Loss of Milton and Miro causes MT curling phenotypes

Next, we focussed on the transport functions of Khc. For example, Pat1 (Protein interacting with APP tail-1) had been shown to link Khc to non-vesicular transport in *Drosophila* oocytes (Fig.3Bii; Loiseau et al., 2010). It is also strongly expressed in the *Drosophila* nervous system (flybase.org: FBgn0029878) but potential neuronal cargoes are unknown. Pat1 function can be eliminated by the gene-specific small deficiencies *Pat1^robin^* and *Pat1^grive^* which represent viable null alleles (Loiseau et al., 2010). When analysing cultures of larval neurons homozygous for either allele, we did not find any obvious enhancement of MT curling (Fig.3Cii).

The linker protein Milton and its binding partner Miro (a small GTPase) link the C-terminus of Khc to organelles including mitochondria (Fig.3A,Biii; Harbauer, 2017; Misgeld and Schwarz, 2017; Sheng, 2017; Smith and Gallo, 2018) and potentially peroxisomes (Castro et al., 2018; Covill-Cooke et al., 2017; Okumoto et al., 2018; Tang, 2018).

Using the loss-of-function mutant alleles *milt^92^*, *Miro^Sd32^* or *Miro^B682^*, we first confirmed the functional contributions of Milt and Miro in primary fly neurons. For this, we stained homozygous mutant neurons with mitoTracker and anti-Syt. We found the axonal localisation of Syt to be unaffected whereas mitochondria were strongly reduced in number, thus confirming the expected cargo specificity (Fig.2E-G,L-P). The mitochondrial phenotype was milder for loss of Miro than Milt, as is consistent also with previous findings in fly neurons *in vivo* as well as mouse neurons (Glater et al., 2006; Guo et al., 2005; López-Doménech et al., 2018; Russo et al., 2009; Vagnoni et al., 2016). In *milt^92^* mutant neurons, mitochondria were virtually absent from axons, restricted mostly to cell bodies and proximal axon segments (Fig.2L). This absence of mitochondria as the major ATP source does not affect synaptic transport because it is self-sufficient through local glycolysis on transported vesicles (Fig.S3A,B; Hinckelmann et al., 2016; Zala et al., 2013).

Having confirmed that the *milt^92^, Miro^Sd32^* or *Miro^B682^* alleles selectively inhibit mitochondrial transport, we then assessed potential impacts on MT organisation. We found that all three mutant alleles caused significant increases in MT curling (Fig.3Ciii). This finding suggests that loss of Khc-mediated organelle transport triggers MT curling (Fig.3Biii).

### Excessive ROS triggers MT disorganisation

Reduced numbers of axonal mitochondria can be expected to impair local homeostasis of calcium, ATP, reactive oxygen species (ROS), and AAA+ protease-mediated protein quality control systems (Glynn, 2017; Misgeld and Schwarz, 2017; Paupe and Prudent, 2018). We started by manipulating the ROS homeostasis through the application of DEM (diethyl maleate). DEM is an effective inhibitor of the anti-oxidant compound glutathione, hence causing the elevation of ROS levels (Fig.4A; Albano et al., 2015; Dasgupta et al., 2012; Pompella et al., 2003). We found that 12 hr-long application of 100 μM DEM before fixation (from 4.5 to 5DIV) induced robust MT curling (Fig.4F). To validate this finding, we then used genetic tools to generate loss- or gain-of-function conditions for ROS-regulating enzymes (yellow highlighted in Fig.4A):

**Fig.4.**
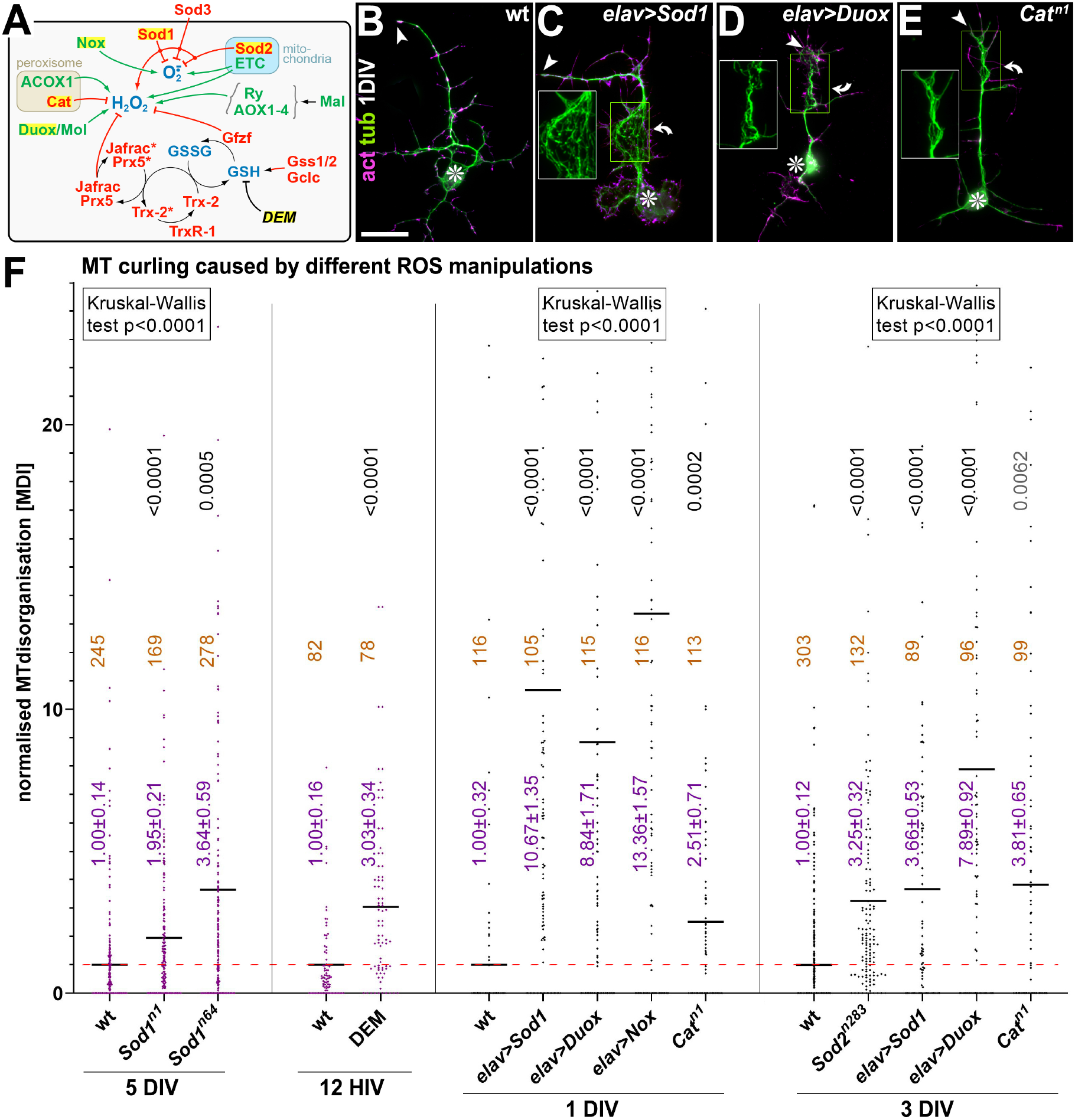
ROS enhancing manipulations cause MT curling phenotypes. **A**) Scheme illustrating the complexity of ROS-regulating systems in *Drosophila*; <u>ROS-generating factors (bold green)</u>: two cytoplasmic NADPH oxidases (Nox/NADPH Oxidase, Duox/Dual oxidase with its essential maturation factor Mol/Moladietz; Khan et al., 2017); enzymes of the mitochondrial EMT/electron transport chain (Wong et al., 2017; Zorov et al., 2014); peroxisomal ACOX1/acyl-CoA oxidase 1 (Walker et al., 2018); Xanthine/aldehyde oxidases (Rosy, AOX1, AOX2, AOX3, AOX4; all jointly silenced by loss of Mal/Maroon-like sulfurtransferase; Marelja et al., 2014); <u>ROS removal mechanisms (red)</u>: superoxide dismutases turn superoxide (O_2_●^−^) into H_2_O_2_ (cytoplasmic CuZn-dependent Sod1, mitochondrial Mn-dependent Sod2, extracellular Sod3); H_2_O_2_ is scavenged by peroxisomal Cat/Catalase and neuronal peroxiredoxins (Jafrac1, Prx5; Cao and Lindsay, 2017; Orr et al., 2013; Smith et al., 2019; Stapper and Jahn, 2018) and the GSH transferase Gfzf (GST-containing FLYWCH zinc-finger protein; Smith et al., 2019; Stapper and Jahn, 2018); the latter three depend on the redox cycle of the Glu-Cys-Gly tripeptide GSH/Glutathione, synthesised by glutathione synthetases (Gss1, Gss2) and Gclc/Glutamate-cysteine ligase (Smith et al., 2019; Stapper and Jahn, 2018) and regenerated via Thioredoxins (primarily Trx-2 in neurons; Orr et al., 2013; Tsuda et al., 2010) and Thioredoxin reductases (primarily TrxR-1 in neurons; Orr et al., 2013; Smith et al., 2019); <u>pharmacological agents (black italics)</u>: DEM/diethyl maleate blocks the GSH system (Pompella et al., 2003); agents/factors used in our study are highlighted in yellow. **B-E**) Examples of neurons, either wild-type (wt) expressing Sod1 or Duox (driven by *elav-Gal4*) or homozygous for *Cat^n1^*, all cultured for 1DIV and stained for actin (act, magenta) and tubulin (tub; green); asterisks indicate cell bodies, arrow heads axon tips, curved arrows areas of MT curling; yellow emboxed areas are shown as 1.5-fold enlarged insets (green channel only); scale bar in B represents 20μm in B-E. **F**) Quantification of MT curling phenotypes measured as MT disorganisation index (MDI) and normalised to wild-type controls (red stippled line); bars at bottom indicate type of culture (‘1/3/5 DIV/HIV’, embryonic neurons 1/3/5 days/hours *in vitro*); mean ± SEM is indicated in blue, numbers of analysed neurons in orange, results of Mann Whitney rank sum tests are shown in grey/black.

Firstly, we used a null allele of Catalase (*Cat^n1^*), an enzyme removing hydrogen-peroxide (H_2_O_2_; Walker et al., 2018), and null alleles of two members of the Superoxide dismutase family, Sod1 (*Sod1^n1^* or *Sod1^n64^*) and Sod2 (*Sod1^n283^*), which convert superoxide anions (O_2_^−^) into H_2_O_2_ (Palma et al., 2020). Of these, Catalase is enriched in peroxisomes, copper-zink-dependent Sod1 (linked to amyotrophic lateral sclerosis/ALS1; #105400; Saccon et al., 2013) is primarily cytoplasmic, and the manganese-dependent Sod2 enzyme is predominantly mitochondrial (Fig.4A). When assessed in primary *Drosophila* neurons, the functional deficiencies of either Sod1, Sod2 or Cat caused MT curling (Fig.4F).

Secondly, we used targeted expression (a) of *Nox* (NADPH oxidase) to enhance O_2_^−^ levels, (b) of *Sod1* to reduce O_2_^−^ levels and enhance H_2_O_2_, and (c) of *Duox* (Dual oxidase) to increase H_2_O_2_ levels (Anh et al., 2011; Bedard and Krause, 2007; Zelko et al., 2002). All these manipulations caused increased MT curling (Fig.4F).

Taken together these results clearly indicate that insults to ROS homeostasis have a strong tendency to trigger MT curling. Upregulation of either O_2_^−^ or H_2_O_2_ seems to cause this effect, although O_2_^−^ may elicit its effects through conversion into H_2_O_2_ (Bedard and Krause, 2007).

### Harmful ROS appears to relate to mitochondria and links loss of Khc or Milt to MT curling

To assess whether harmful ROS might be responsible for linking loss of Khc/Milton/Miro to MT curling, we treated mutant neurons with Trolox (6-hydroxy-2,5,7,8-tetramethylchroman-2-carboxylic acid), an α-tocopherol/vitamin E analogue that displays beneficial antioxidant effects in many cell systems by inhibiting fatty acid peroxidation and quenching singlet oxygen and superoxide (Giordano et al., 2020). Neurons mutant for *Khc^8^* or *milt^92^* were either pre-cultured for a day in the presence of 100μM Trolox, or they were cultured directly for 5 days with Trolox. Under both conditions, application of Trolox strongly suppressed or even abolished the MT curling phenotype of *Khc^8^* and *milt^92^* mutant neurons (Fig.5); this indicated harmful ROS to be the main reason for MT bundle disintegration. It might explain why rat neurons depleted of the Khc homologue KIF5C were reported to be more vulnerable to H_2_O_2_ application (Iworima et al., 2016).

**Fig.5.**
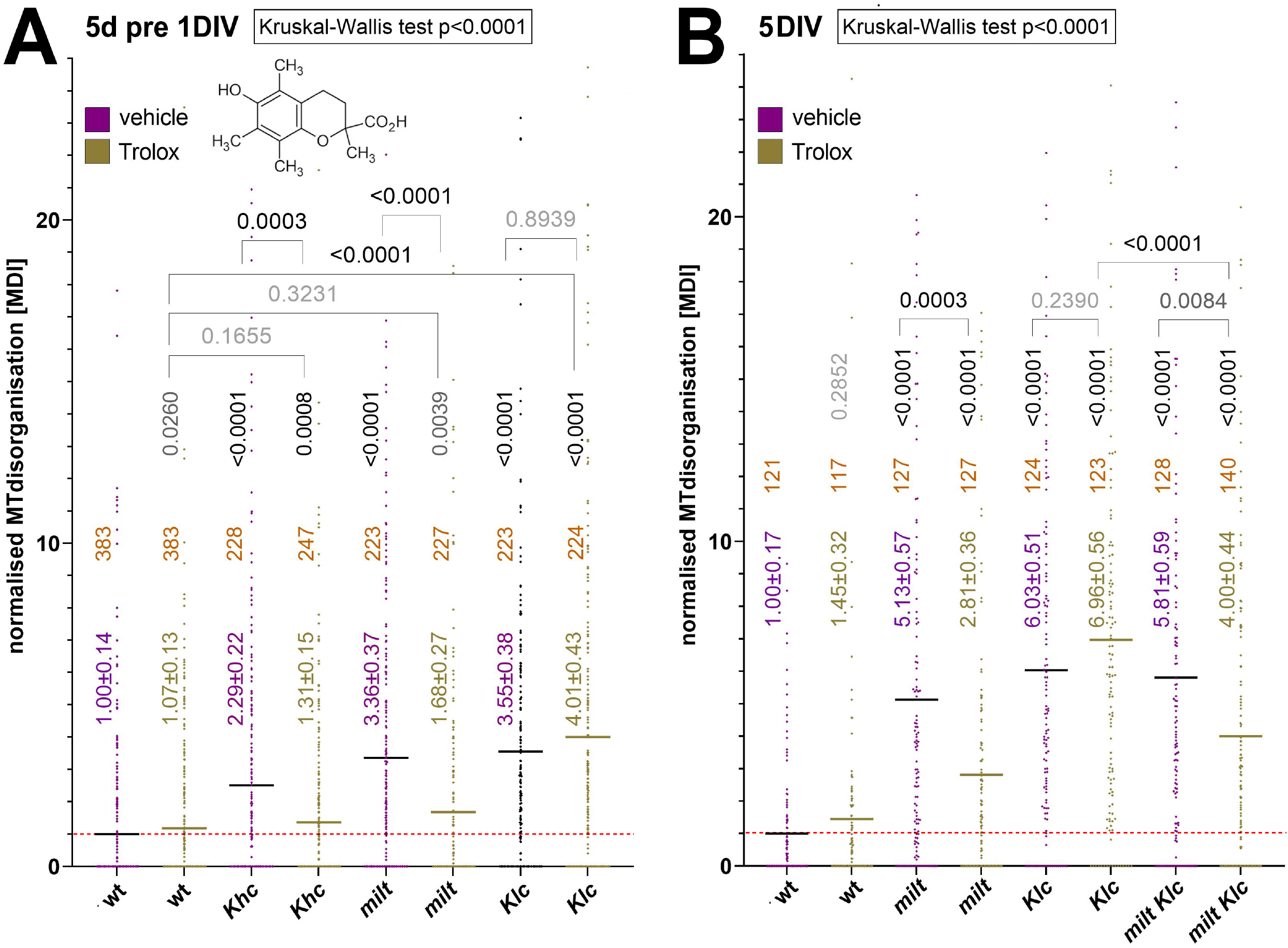
Ameliorating effects of Trolox on mutant MT curling phenotypes. Quantification of MT curling phenotypes measured as MT disorganisation index (MDI) and normalised to wild-type controls (red stippled line); neurons of different genotype (indicated below) were cultured for 1day after preculture (A) of for 5 days (B) in the presence of vehicle (blue) or 100μm Trolox (green; molecule depicted in A); mean ± SEM is indicated in blue/green, numbers of analysed neurons in orange, results of Mann Whitney rank sum tests are shown in grey/black.

To understand how loss of Khc and Milt might trigger harmful levels of ROS, we first set out to identify the potential source. For example, loss of Catalase causes MT curling (Fig.4A,F), potentially suggesting that peroxisomes are required to keep H_2_O_2_ levels down. To test this possibility, we blocked peroxisome biogenesis using the *Pex3^2^* mutation (Faust et al., 2014). Abolishing peroxisomes in this way did not induce any obvious MT curling phenotypes, but the *Pex3^2^* mutant neurons had shorter axons (Fig.S4) potentially due to lack of peroxisomal lipidogenesis (Wanders et al., 2020). The absence of obvious MT phenotypes seems to contradict MT curling observed upon Catalase deficiency (Fig.4F), but it might be explained by observations that Catalase can localise outside peroxisomes in the cytoplasm (Zhou and Kang, 2000).

We concluded that MT curling observed upon loss of Khc, Milt or Miro is more likely to link to their roles in axonal transport of mitochondria; disturbing mitochondrial dynamics could either generate harmful ROS (via the ETC; Fig.4A) or affect their ability to quench local ROS (via Sod2; Fig.4A).

### ROS-absorbing properties rather than disrupted fission/fusion of mitochondria might provide links to MT curling

Loss of Khc/Milt/Miro might cause harmful ROS by affecting fission/fusion processes required to maintain a healthy mitochondrial population (Cagalinec et al., 2013; Liu et al., 2009; Wang et al., 2015). In support of this notion fission/fusion factors are linked to axonopathies; this is the case for the fission factor DNM1L/DYNAMIN-LIKE PROTEIN 1 (Optic atrophy 5; OMIM^®^ #610708), as well as the fusion factors OPA1/OPA1 MITOCHONDRIAL DYNAMIN-LIKE GTPase (Optic atrophy 1; #165500) and MFN/MITOFUSIN (CMT2A2A, CMT2A2B, HMSN6A; #609260, 617087, 601152).

To test whether loss of fission/fusion is a condition that affects MT bundling, we used mutant alleles abolishing the functions of the fly homologues of mammalian DNM1L (*Drp1^T26^*; Dynamin related protein 1), of mammalian OPA1 (*Opa1^s3475^*; Optic atrophy 1) and of mammalian MFN (*Marf^B^*; Mitochondrial assembly regulatory factor). In axons of wild-type neurons, mitochondria mostly displayed dash-like shapes (Fig.S5A) and occasionally appeared dot-like or formed longer lines (not shown). In contrast, within axons of neurons with impaired fusion (*Opa1^s3475^* or *Marf^B^*) mitochondrial shapes were primarily short and dot-like (Fig.S5C,D), whereas loss of fission (*Drp1^T26^*) caused string-of-pearl arrangements where a continuous thread of mitochondria ran all along the main axon but was excluded from side branches (Fig.S5B). These findings are consistent with reports for mammalian neurons (Smirnova et al., 2001; Uo et al., 2009; Yu et al., 2011).

When analysed for MT organisation, none of the three fission/fusion-deficient conditions caused curling in axons, neither at 5 DIV nor upon pre-culture (Fig.S5E,F). This might suggest that MT curling upon loss of Khc/Milt/Miro is unlikely to be caused by mitochondrial fission/fusion defects; the absence of fission/fusion events seems not to affect mitochondria in ways that cause harmful ROS leakage, as is also consistent with views of other authors (Misgeld and Schwarz, 2017). Accordingly, also MFN2-deficient mouse neurons seem not to experience oxidative stress (Baloh et al., 2007). A further argument against the involvement of fission/fusion is based on the observation that mitochondria in Milt-deficient neurons tend to stay in the cell body: it is unlikely that harmful ROS generated in the soma were to reach the distal axon via long-range diffusion, especially when considering the abundance of ROS-buffering systems (Fundu et al., 2019; Kükürt et al., 2021; Oswald et al., 2018).

We prefer therefore the explanation that MT curling in Khc/Milt/Miro-deficient neurons (Fig.2P) might be caused by the absence of mitochondria from critical positions in the axon, thus depleting these areas from Sod2 activity (for details see Discussion).

### Klc also causes MT curling but through a mechanism distinct from Milt

Given the comparable strength of MT phenotypes upon loss of Khc, Milt or Miro (Fig.3Ciii) and their shared links to harmful ROS production (Fig.5), mitochondrial transport defects seemed to offer the perfect explanation for why loss of Khc induces MT curling. We expected therefore that loss of the vesicular transport linker Klc would not cause these MT phenotypes.

Surprisingly, we found that also loss of Klc (*Klc^8ex94^* or *Klc^8ex94/Df^* mutant neurons at 5 DIV) caused MT curling, and that this phenotype was at least as strong as observed with Khc or Milt deficiency (Fig.3Ciii,iv). We obtained the same results when performing 5d pre-cultures with *Khc^8^*, *Klc^8ex94^* and *milt^92^* mutant neurons (to deplete their maternal products; Figs.6A-E), confirming that the three factors generate comparably strong MT curling phenotypes.

**Fig.6.**
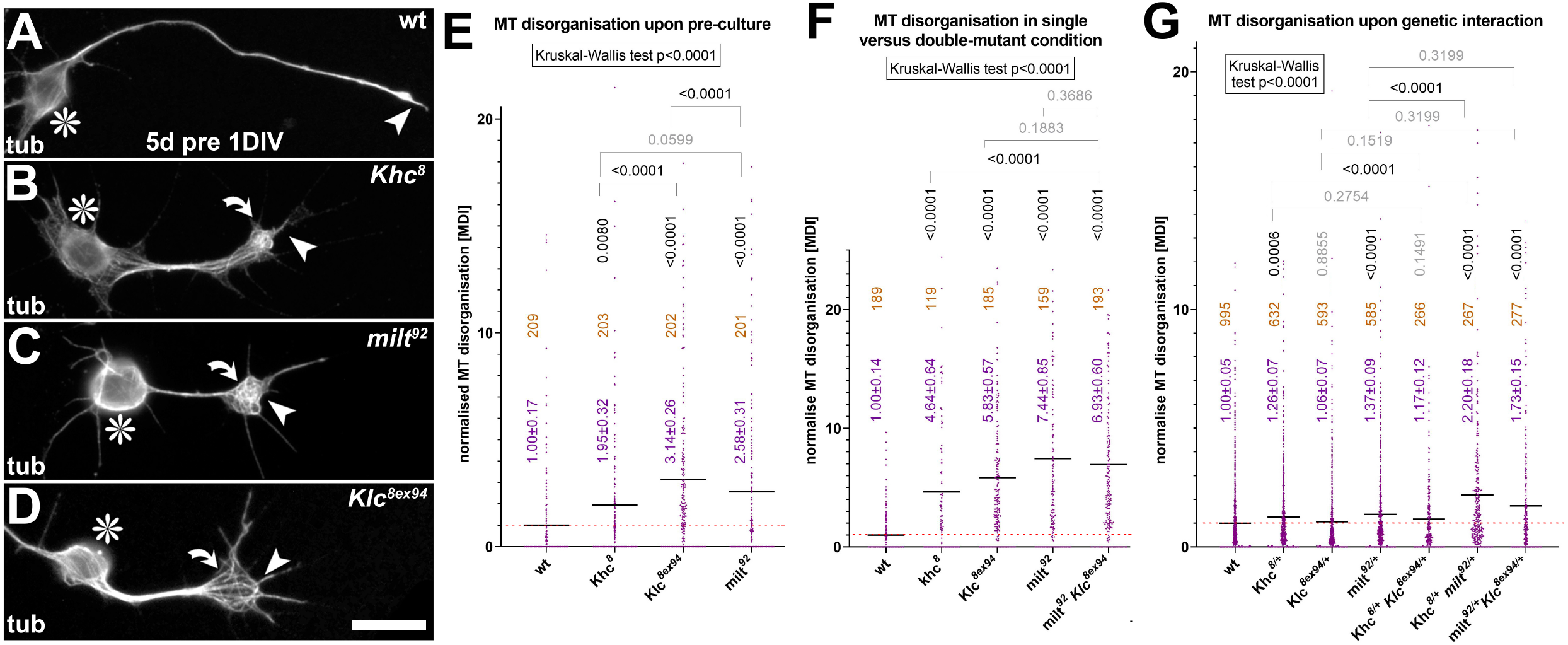
Genetic studies of functional links between Khc, Milt and Klc. **A-D**) Examples of neurons, either wild-type (wt) or homozygous for *Khc, milt* or *Klc* null mutant alleles, cultured for 1DIV following 5d pre-culture (to deplete maternal product) and stained against tubulin (tub); asterisks indicate cell bodies, arrow heads axon tips, curved arrows areas of MT curling; scale bar in D represents 20μm in A-D. **E-G**) Quantification of MT curling phenotypes measured as MT disorganisation index (MDI) and normalised to wild-type controls (red stippled line) shown for precultured neurons (E; as in A-D), single/double-homozygous mutant neurons (F) and upon genetic interaction (G; single heterozygous and trans-heterozygous); mean ± SEM is indicated in blue, numbers of analysed neurons in orange, results of Mann Whitney rank sum tests are shown in grey/black.

To establish whether Klc might work synergistically with Khc and Milt, we applied Trolox to *Klc^8ex94^* mutant neurons. However, in contrast to *Khc^8^* and *milt^92^* mutant neurons, the MT curling in Klc-deficient neurons was not suppressed by Trolox, neither in pre-cultured neurons nor in 5DIV cultures (Fig.5). This clearly demonstrated that Klc works through an independent mechanism.

### Klc’s links to MT curling do not depend on vesicular cargo transport

We first tested whether Klc’s impact on MT regulation might link to vesicular cargo transport, capitalising on reports that Khc-mediated vesicular cargo transport requires a protein complex of a number of factors including Klc and Sunday driver (Syd, the JIP3 homologue; Fig.3Biv; Gindhart et al., 2003; Horiuchi et al., 2005; Koushika, 2008). Accordingly, functional loss of either Klc or Syd was shown to abolish Khc-mediated vesicular transport, with motoraxons in peripheral larval nerves displaying synaptic protein accumulation that were similarly strong upon Klc or Syd deficiency as observed upon loss of Khc (Bowman et al., 2000; Füger et al., 2012; Gauger and Goldstein, 1993; Gindhart et al., 1998; Hurd and Saxton, 1996; Pilling et al., 2006). Equally in primary culture, we found that the number of Synaptotagmin-positive dots in axons was reduced in *Klc^8ex94^* and *syd^z4^* null mutant neurons, and the phenotypes were similarly strong as observed in Khc-deficient neurons (Fig.2B,G,O). These effects seem specific to vesicular cargo transport since mitochondrial numbers in axons of *Klc^8ex94^* mutant neurons appeared normal (Fig.2N,P).

Our data confirm therefore that Klc, Syd and Khc closely co-operate during vesicular cargo transport in primary *Drosophila* neurons. However, in contrast to severe MT curling in Klc- and Khc-deficient neurons, *syd^z4^* or *syd^z4/Df^* mutant neurons at 5DIV failed to show similar phenotypes (Fig.3Civ). This observation was further supported by experiments where we knocked down GAPDH (glyceraldehyde-3-phosphate dehydrogenase), an enzyme required for glycolysis that is known to fuel the axonal transport of vesicles but not of mitochondria (details in Fig.S3A,B; Zala et al., 2013). When GAPDH was knocked down in primary *Drosophila* neurons, axons displayed a reduction in synaptic dots but not in mitochondrial numbers (Fig.S3C), as is consistent with analyses in larval nerves (Zala et al., 2013). However, GAPDH knock-down did not cause obvious MT curling phenotypes (Fig.S3C), thus mirroring results obtained with Syd deficiency.

We concluded that blocking vesicular axonal transport appears not to be a cause for MT curling, and that loss of Klc is likely to trigger its MT phenotypes through a different mechanism. This view was also supported by genetic interaction studies using trans-heterozygous pairings of *Khc^8^*, *milt^92^* and *Klc^8ex94^* (i.e. combining heterozygosity for two genes at a time in the same neurons). Of the three constellations, only *Khc^8/+^ milt^92/+^* trans-heterozygote mutant neurons generated a MT curling phenotype that was significantly enhanced over single heterozygous conditions (Fig.6G), supporting functional links between Khc and Milt but not with Klc.

Taken together, MT curling upon loss of Khc appears to relate to Milt/Miro-mediated mitochondrial transport as explained before, but not to Klc-mediated vesicular transport. Milt, Miro and Khc seem to have comparably strong mutant phenotypes because their loss leads to the same transport defect, whereas phenotypes observed upon loss of Klc (which has binding sites on Khc that overlap with those of Milt; details in Fig.3A) seem not to relate to its function as a transport linker but work through an entirely different mechanism.

### Excessive pools of active Khc might explain the Klc-deficient MT phenotype

We hypothesised that MT curling upon loss of Klc may relate to its roles in regulating the activation state of Khc. Thus, Khc pools that are not linked to cargo tend to be auto-inhibited and detached from MTs. This inactivation requires intramolecular loop formation via binding of the N-to the C-terminus, and this also involves the association with Klc (co-regulated through its own auto-inhibition/activation mechanism; Figs.3A,Biv; Bowman et al., 2000; Koushika, 2008; Verhey and Hammond, 2009; Verhey et al., 1998; Wong and Rice, 2010; Yip et al., 2016). In non-neuronal cells, overriding auto-inhibition of the Khc-Klc complex causes MT curling (Paul et al., 2020; Randall et al., 2017).

To test whether Klc-deficient MT curling in neurons might involve excessive pools of active Khc, we first targeted the expression of GFP-tagged constructs of Khc to neurons. We found that full-length Khc::GFP was homogeneously distributed along axons and failed to increase MT curling when analysed at 5DIV (Fig.7), consistent with the idea that extra pools of Khc tend to be inactive and detached from MTs.

**Fig.7.**
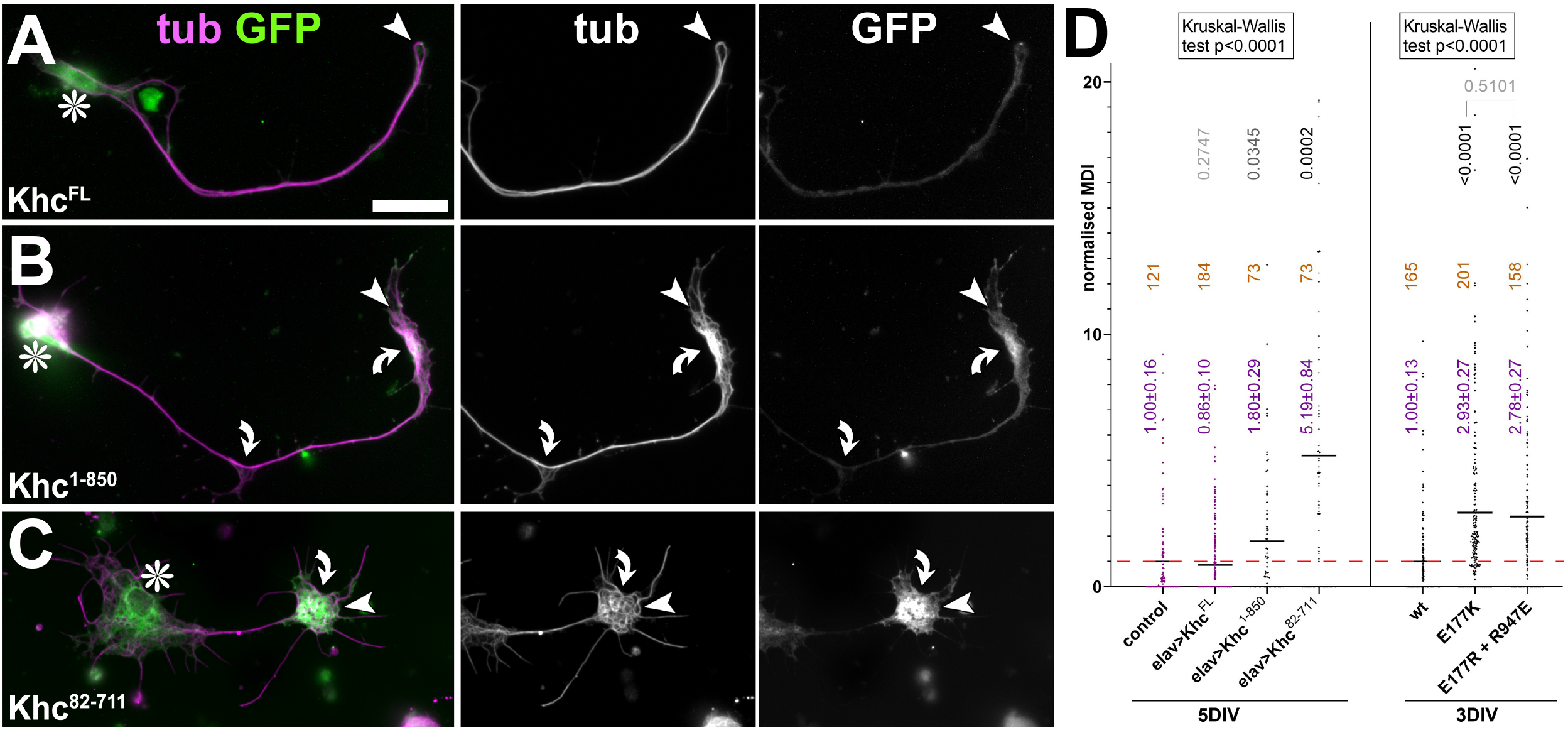
Impacts of activated Khc on MT curling. **A-C**) Examples of neurons at 5DIV expressing different Khc constructs (indicated bottom left; compare Fig.3A) and stained for tubulin (tub, magenta) and GFP (green), also shown as greyscale single channel images on the right; asterisks indicate cell bodies, arrow heads axon tips, curved arrows areas of MT curling; scale bar in A represents 20μm in all images. **D**) Quantification of MT curling phenotypes measured as MT disorganisation index (MDI) and normalised to wild-type controls (red stippled line); genotypes are shown below, also indicating the culture period (5DIV, 3DIV); mean ± SEM is indicated in blue, numbers of analysed neurons in orange, results of Mann Whitney rank sum tests are shown in grey/black.

We then expressed two non-inactivating Khc derivatives (Khc^1-811^::GFP and Khc^82-711^::GFP; top of Fig.3A) which both lack the C-terminal domain needed for auto-inhibition but also for their roles in cargo transport and MT sliding (dark grey and green in Fig.3A). When analysed in neurons at 5DIV, both constructs accumulated at axon tips, as is typical of non-inactivating kinesins (Niwa et al., 2013). Of these, Khc^1-850^::GFP caused a very mild MT phenotype, suggesting that extra pools of free-running Khc *per se* cause little harm (Fig.7B,D). In contrast, Khc^82-711^::GFP caused severe MT curling (Fig.7C,D), potentially because this truncated form also has a small N-terminal deletion – and short deletions of the N-terminus have been shown to display damaging effects on MTs (Budaitis et al., 2021; details in Fig.3A). In principle, our findings with this dys-regulated construct supported our previously published hypothesis that the activity of kinesins is harmful to axonal MT bundles and can explain MT curling (Hahn et al., 2019; Prokop, 2021).

However, since all essential C-terminal binding sites were removed in Khc^1-850^::GFP and Khc^82-711^::GFP (Fig.3A), our experiments so far only assessed free-running Khc that could not engage in movement of any cargo (Fig.3A,B). We hypothesised that active transport would be expected to generate higher forces than free-running Khc and, hence, be more challenging to MT bundles. We therefore tested the genomically engineered *Khc^E177K^* and *Khc^177R, R947E^* mutant alleles (Fig.3A), which have point mutations in the E177 and R947 residues that are known to form a required salt bridge with each other during auto-inactivation (Kaan et al., 2011; Kelliher et al., 2018); these mutant alleles cause lethality and distal accumulations of Khc (Brendza et al., 1999; Kelliher et al., 2018). When analysing *Khc^E177K^* and *Khc^E177R, R947E^* homozygous mutant neurons at 3DIV, we found that they display robust MT curling (Fig.7D). This clearly indicated that extra pools of actively engaging Khc harm MT bundles, which might therefore explain the *Klc* mutant phenotype (see Discussion).

## Discussion

### Using MT bundles of *Drosophila* neurons as a powerful approach to dissect motor-related patho-mechanisms

Motor proteins involved in axonal transport clearly are key drivers of neuronal survival, yet their links to axonopathies remain poorly understood and speculative (Coleman, 2005; Guo et al., 2020; Kawaguchi, 2013; Sleigh et al., 2019). Here, we aimed to unravel concrete mechanisms through which motor protein loss can affect axons.

Our approach was unprecedented in that we performed a systematic genetic study in one standardised neuron system and used MT bundles as key readout. We studied the organisation of axonal MT bundles because they are good indicators of axon integrity (Prokop, 2020) that are easy to quantify and have an intricate interdependent relationship with motor proteins (Prokop et al., 2013). The key phenotype we observed upon motor manipulation is MT curling which appears conserved across species, since depleting Dynein or the kinesin-1 linker JIP3 causes the same kind of curling in axons of mammalian neurons (Ahmad et al., 2006; Rafiq et al., 2020).

The easily accessible and quantifiable MT curling readout allowed us to determine roles of motor proteins and their interactors, or of proteins regulating potential downstream processes - and many of these factors have known links to neurodegeneration or axonopathies. Our approach was facilitated by using a standardised neuronal culture system in which findings could be integrated and were highly accessible to powerful *Drosophila* genetics. The additional advantage of this system is low genetic redundancy of factors involved in axonal transport, with one *Drosophila* gene having on average almost 3 mammalian orthologues (Khc/Kif5: 1 paralogue in fly *vs*. 3 paralogues in mammals; Miro/RHOT: 1 *vs*. 2; Milt/TRAK: 1 *vs*. 2; Klc: 1 *vs*. 4; Marf/MFN: 1 *vs*. 2; Unc-104/Kif1: 1 *vs*. 3). This low redundancy in fly enormously facilitates loss-of-function analyses and combinatorial genetics.

Capitalising on these advantages, our unprecedented strategy enabled us to generate new understanding and conceptual explanations. So far, we found that deficiencies of three motor proteins (Khc, Unc-104, Dhc) cause MT curling. Notably, the homologues of all three factors have OMIM^®^-listed links to human axonopathies (see Introduction) potentially reflecting evolutionarily conserved mechanisms of axon pathology that might even be shared between these motor protein classes.

For example, we found that phenotypes upon loss of Khc and Unc-104 are very similar with respect to enhanced MT curling and the reduction in axonal numbers of mitochondria and synaptic dots (Figs.1, 2), and both seem involved in mRNA transport (L.M.P.C., unpublished results; Lyons et al., 2009). This functional overlap is in agreement with reports that kinesin-1 and −3 collaborate during transport (Arpağ et al., 2019; Zahavi et al., 2021). Consequently, these two motors might therefore link to axonopathy through comparable mechanisms.

### An intricate relationship: kinesins simultaneously harm and care for MT bundles

The challenges of studying axonal transport are ample due to (1) the parallel involvement of different motor protein classes (which might act redundantly; Hirokawa et al., 2010; see previous section), (2) the enormous wealth of their cargoes, (3) the involvement of many different linkers (that might interact promiscuously with different motors; Brady and Morfini, 2017; Drerup et al., 2016; Gindhart, 2006; Hirokawa et al., 2010; Maday et al., 2014), (4) additional roles in slow transport (transient ‘hitchhiking’ of proteins on transported vesicles; Roy, 2020; Tang et al., 2013), and (5) complications caused by the interdependence of kinesins and dynein/Dynactin (Hancock, 2014; Moughamian et al., 2013; Twelvetrees et al., 2016; potentially explaining the rather counter-intuitive observation that loss of anterograde Khc transport causes distal ER accumulations; details in legend of Fig.S1).

For kinesin-1 alone (Fig.3B), we tested roles of Khc in MT sliding (*Khc^mutA^*), roles of Khc/Milt/Miro in mitochondrial/peroxisomal transport, of Khc/Klc/Syd/GAPDH in vesicular transport, of Khc/Pat1 in potential non-vesicular transport, and potential roles of Klc and certain Khc domains/residues in Khc auto-inhibition. These extensive studies still left out further known linkers, such as SKIP/SNW1/SKIIP and Arl8 (lysosome transport; Keren-Kaplan and Bonifacino, 2021; Rosa-Ferreira and Munro, 2011; Rosa-Ferreira et al., 2018) or Tropomyosin (mRNA transport; Fig.3A; Dimitrova-Paternoga et al., 2021; Veeranan-Karmegam et al., 2016). Nevertheless, the analyses we performed suggested two distinct mechanisms:

Firstly, Khc/Milt/Miro-mediated transport is required to uphold ROS homeostasis, with harmful ROS being a strong inducer of MT curling (demonstrated by our studies with DEM, Trolox and ROS-regulating enzymes; Figs.4, 5). Secondly, we found that the movement and active transport of Khc along MTs damages axonal bundles, as demonstrated by the expression of Khc deletion constructs and analyses of non-inactivating *Khc* mutant alleles (Fig.7). These latter findings align with published *in vitro* experiments demonstrating kinesin-1-induced MT damage (Andreu-Carbó et al., 2021; Budaitis et al., 2021; Dumont et al., 2015; Triclin et al., 2021; VanDelinder et al., 2016), MT curling observed in kinesin-1-based gliding assays *in vitro* (Hahn et al., 2019; Lam et al., 2016), and the curling observed upon kinesin-1 activation in non-neuronal cells (Paul et al., 2020; Randall et al., 2017). Notably, MT curling is not specifically linked to motor proteins, but is similarly observed upon loss of various MT-binding and -regulating proteins (Hahn et al., 2019) and in a model of chemotherapy-induced peripheral neuropathy (Rozario et al., 2021).

All these causes of MT curling, including the two mechanisms described in this work, can be explained with the previously proposed “local axon homeostasis” model (Hahn et al., 2019) and the subsequently derived “dependency cycle of axon homeostasis” (Prokop, 2021; details in Fig.8). These models propose that kinesins that trail along MT bundles during axonal transport (‘2’ in Fig.8) pose a mechanical challenge that leads to MT curling (‘3’); active machinery of MT-regulating proteins and support through the cortical actin-spectrin sleeve is therefore required to prevent disintegration and maintain these bundles long-term (‘4’). However, the machinery that maintains MT bundles is itself dependent on materials and physiology provided by axonal transport (‘5’), thus establishing a cycle of mutual dependency where interruption at any point will have a knock-on effect on all other aspects of axon physiology and function (Prokop, 2021). The mechanisms we described here act in either direction of this cycle (thick read arrows in Fig.8).

**Fig.8.**
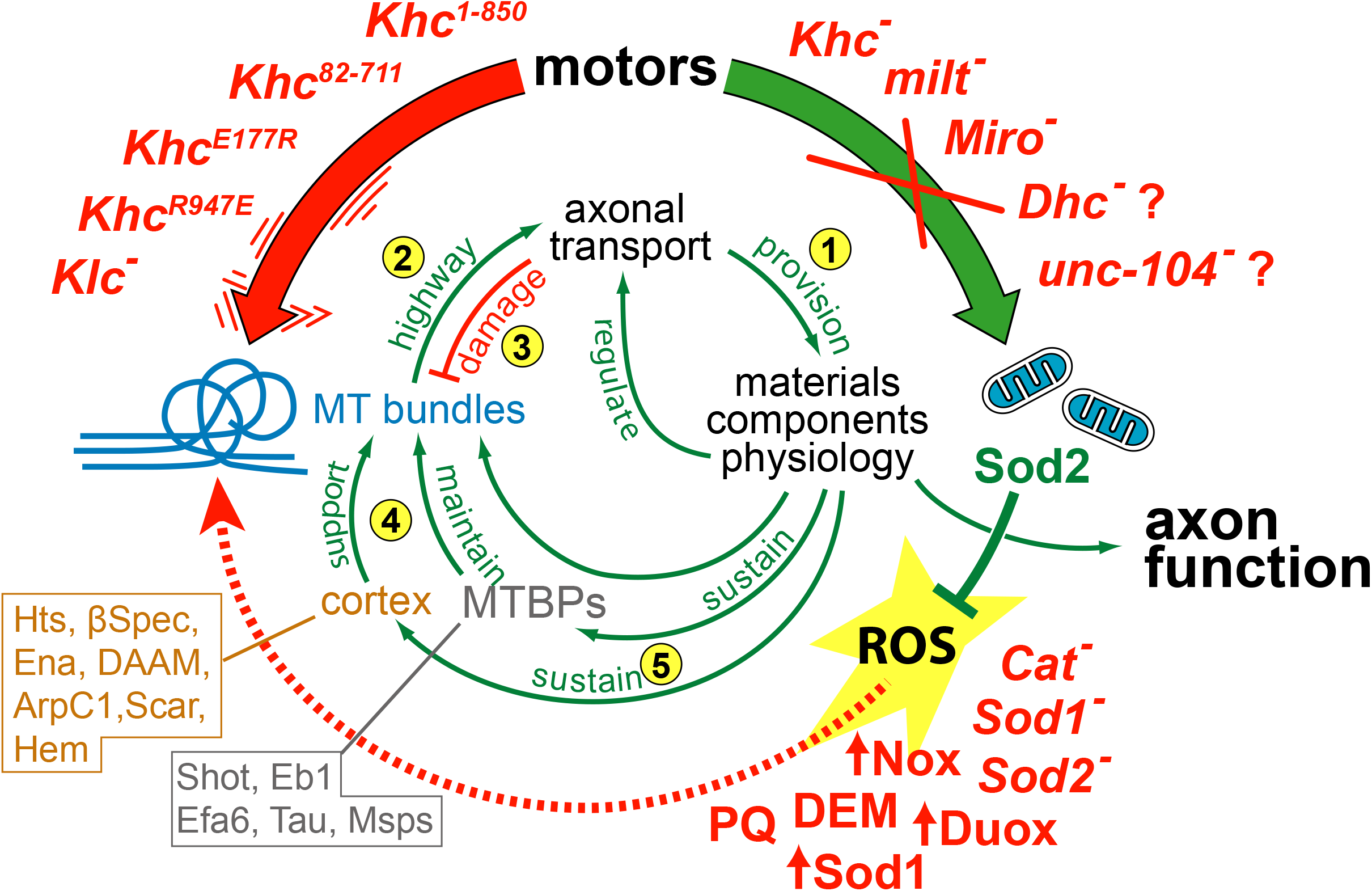
Mapping findings on the dependency cycle of local axon homeostasis. The numbered green arrows and red T-bar make up the previously published ‘dependency cycle of local axon homeostasis’ (Prokop, 2021): 1) axonal transport provides materials, components and organelles required for axon function; 2) this transport requires MT bundles as the essential highways; 3) however, this live-sustaining transport damages MT bundles; 4) the axonal cortex and MT binding proteins (MTBPs) support and maintain MT bundles (emboxed names in orange and grey at bottom left list factors that were shown in the *Drosophila* neuron culture system to be involved in bundle-maintaining cortical and MT regulation;Alves-Silva et al., 2012; Hahn et al., 2021; Qu et al., 2019; Qu et al., 2017); 5) bundle maintenance requires transport-dependent components and physiology, thus closing the circle. The original model of ‘local axon homeostasis’ comprised arrows 1-4 (Hahn et al., 2019). Khc contributes to the MT bundle damage, and this is enhanced by non-inactivating mutations (vibrating red arrow and *Khc* alleles top left). Loss of function of Khc, Milt, Miro, Unc-104 and Dhc contribute to mitochondrial transport (large green arrow, top right). Mitochondria harbour Sod2 that can quench harmful ROS (green T-bar); 8 independent pharmacological and genetic manipulation of ROS regulation (bottom right) demonstrated that dysregulation of ROS causes MT curling (dashed red arrow). Examples of mammalian factors that can be mapped onto this cycle are explained in the Discussion and previous reviews (Hahn et al., 2019; Prokop, 2021).

### Mitochondria regulate ROS homeostasis required for MT bundle maintenance

Our finding that harmful ROS is a key trigger of MT curling aligns with reports that actin as well as MTs are modified or even damaged by ROS (Goldblum et al., 2021; Wilson et al., 2016; Wioland et al., 2021) and that oxidative stress induces axon swellings in models of Parkinson’s disease, multiple sclerosis or ALS (Czaniecki et al., 2019; Nikić et al., 2011; Song et al., 2013). Unfortunately, a more generalised statement cannot be made because axonal MTs have rarely been analysed in oxidative stress experiments (De Vos et al., 2007; Debattisti et al., 2017; Fischer et al., 2012; Saccon et al., 2013; Song et al., 2013).

Pinpointing the precise source of harmful ROS upon Khc/Milt/Miro loss is a tedious task when considering (1) the intricate network of ROS regulation (Fig.4A) where manipulations of very different regulators caused comparable phenotypes (Fig.4F), and (2) the spectrum of organelles involved in ROS homeostasis regulation: these involve the finely tuned mitochondria-peroxisome system (Fransen et al., 2017; Pascual-Ahuir et al., 2017), but also the ER which contains oxidases required for protein folding (Hudson et al., 2015). Unfortunately, removing or affecting the ER to assess its involvement is not trivial (O’Sullivan et al., 2012; Yalcin et al., 2017); but our studies of *Pex3* mutant conditions suggested that peroxisomes are unlikely to link to MT curling (Fig.S4). This said, peroxisomes certainly play important further roles in maintaining healthy axons (Wali et al., 2016).

In our view, the most likely organelles involved in MT curling are the mitochondria. We were surprised to find that not the presence of damaged mitochondria leaking harmful ROS seems to trigger MT curling (Fig.S5), but rather the absence of mitochondria. This is best illustrated by *milt* mutant neurons where mitochondria are mostly restricted to cell bodies (Fig.2L,P), yet strong ROS-induced MT curling occurs in axons (Figs.3Ciii, 5, 6E).

As already mentioned in the Results part, we believe that the best model combining all observations is the absence of mitochondria and Sod2 as their ROS scavenger from critical locations in axons. For example, we know from live imaging experiments that MT curling starts at growth cones or branch points (A.V., unpublished data), and both are typical sites where mitochondria localise (Bunge, 1973; Mandal and Drerup, 2019). Failure to quench harmful ROS in these critical locations could therefore promote the initiation of MT curling; this would also explain why drastic mitochondrial depletion upon Milt deficiency triggers similarly strong MT curling as moderate depletion upon loss of Khc or Miro (Figs.2P, 3C): not the number of mitochondria is essential but their adequate localisation, and this aspect is regulated through a Khc/Milt/Miro-dependent mechanism (Misgeld and Schwarz, 2017).

The drastic depletion of axonal mitochondria upon Milt deficiency as compared to the moderate number reductions upon loss of Khc, Unc-104 or Dhc (Fig.2P) might suggest Milt as a ‘master linker’ for mitochondrial transport in fly neurons. Indeed, Milt is known to link to Khc and Dynein in both flies and mammals (Russo et al., 2009; van Spronsen et al., 2013), whereas there are currently no such reports for Unc-104; its mammalian homologue Kif1 was reported so far to perform mitochondrial transport through KBP (Kif1 binding protein; Campbell et al., 2014; Nangaku et al., 1994; Tanaka et al., 2011; Wozniak et al., 2005).

### Khc activation as a further factor leading to MT curling

As discussed above, our data with non-inactivating constructs and mutant alleles of Khc strongly suggested that excess engagement of this motor can trigger MT curling (Fig.7). Since Klc is involved in Khc auto-inhibition, the MT curling phenotype we observe in axons of Klc-deficient neurons might therefore link to this mechanism as a potential cause for axonopathy.

Also Kif1A/Unc-104 undergoes inactivation involving intramolecular loop formation and the KBP linker (Cong et al., 2021; Kevenaar et al., 2016). KBP mutations impair axon growth, are disruptive to axonal MT bundles (Lyons et al., 2008) and cause the devastating neurological disorder Goldberg-Shprintzen syndrome in humans (Chang et al., 2019; Hirst et al., 2017). Similarly, non-inactivating mutations of Kif1A cause spastic paraplegia (Chiba et al., 2019; Gabrych et al., 2019). Also KLC2 has been linked to neuropathy (SPOAN; #609541) and KLC4 mutations have recently been reported to cause excessive axon branching (Haynes et al., 2021). Axonopathy-linked human mutations of Kif5A were mapped exclusively to the motor domain or the very C-terminal end so far, but none were reported in the auto-inactivation domains or the KLC binding site (Nicolas et al., 2018). However, this does not mean that such mutations are not detrimental: mutations affecting auto-inhibition might rather confer lethality (as is the case in *Drosophila*; Brendza et al., 1999; Kelliher et al., 2018) and therefore escape the spectrum of diagnosed diseases.

The modest curling observed upon overexpression of Khc^1-850^::GFP, versus the strong phenotype with genomically engineered *Khc^E177K^* and *Khc^E177E, R947E^* mutant alleles (Fig.3A), suggests that free-running Khc is insufficient to cause a strong phenotype; instead it seems that the C-terminus of Khc has to interact with cargo to generate forces strong enough to affect MT bundles. But what cargoes might be involved?

Some insights might come from *milt^92/92^ Klc^8ex94/8ex94^* double-mutant neurons which show an intriguing pattern: loss of Klc and Milt trigger MT curling through completely different mechanisms (Fig.8), and their phenotypes should therefore be additive. However, double-mutant neurons show the same amount of curling as the single mutants (Figs.5B, 6F). Even more, MT curling in double-mutant neurons is partly cured by Trolox, although Klc-deficiency alone does not respond to Trolox (Fig.5B). The easiest explanation for these findings is that surplus pools of activated Khc triggered by loss of Klc engage in force-generation that depends on mitochondria-derived ATP (unlike vesicular transport; Fig.S3A). Since mitochondria and their ATP are absent from axons of *milt Klc* double-mutant neurons, the extra pool of Khc lacks the necessary fuel to contribute to the joint MT curling phenotype. So far, our attempts to pinpoint such force-generating activities of surplus Khc pools have not been successful: they seem not to involve microtubule sliding and Pat1-mediated transport (Fig.3Bi,ii) as suggested by failed suppression of MT curling in *Khc^mutA/mutA^ Klc^8ex94/8ex94^* or *Pat1^robin/robin^ Klc^8ex94/8ex94^* double-mutant neurons (details in Fig.S6).

### Conclusion

Using our unconventional strategy (MT curling as key readout for systematic genetic analyses in a standardised *Drosophila* primary neuron system) we were able to develop new concepts for how molecular motor mutations might trigger axonopathies. Given the breadth of genetics versus lack of mechanistic detail of our studies, our results are certainly more suggestive than definite. But they are astonishingly consistent with many reports in the field (as mentioned throughout this work) and align well with the ‘dependency cycle of local axon homeostasis’ as a model describing the fundamental principle of axon maintenance and pathology (Prokop, 2021). We hope therefore that our findings and ideas will stimulate further studies in which the model and proposed mechanisms are put to the test, incorporating also other aspects of axon physiology, such as ATP and calcium regulation. Whatever the outcome, such studies will be highly informative and contribute to the battle against a class of diseases that are of enormous socioeconomic burden and personal hardship.

## Conflict of Interest

None of the authors has a conflict of interests

## Acknowledgements

This work was made possible through support by the Biotechnology and Biological Sciences Research Council (BBSRC) to A.P. (BB/I002448/1, BB/P020151/1, BB/L000717/1, BB/M007553/1) and M.L. (BB/R016666/1), by parents to Y.-T.L., H.T. and T.M. The Fly Facility has been supported by funds from The University of Manchester and the Wellcome Trust (087742/Z/08/Z). We thank colleagues and the Bloomington *Drosophila* Stock Center (NIH P40OD018537) for providing fly stocks as mentioned in Materials and Methods.

## Methods

### Fly stocks

All human homology statements are based on information listed on flybase.org (Marygold et al., 2016; Millburn et al., 2016), all statements about genetic links to human diseases on information provided by www.omim.org (Online Mendelian Inheritance in Man^®^; Amberger et al., 2015). The following stocks were used in this study (reference and source provided in brackets; BL indicates Bloomington Drosophila Stock Collection): null mutant alleles (unless indicated differently) we used were

- *unc-104^170^* (Pack-Chung et al., 2007; Tom Schwarz)
- *Klp64D^k1^* (Ray et al., 1999; hypomorphic allele; BL #5578)
- *Klp64D^n123^* (Perez and Steller, 1996; BL #5674)
- *Klp98A^Δ47^* (Derivery et al., 2015; Marcos Gonzalez-Gaitan)
- *Dhc64C^4-19^* (Gepner et al., 1996; BL #5274)
- *Khc^8^* (Saxton et al., 1991; BL #1607)
- *Khc^27^* (Saxton et al., 1991; Isabel Palacios)
- *Klc^1ts^* (Saxton et al., 1991; BL #31994; a temperature-sensitive allele which is homozygous viable at 18°C but usually kept over balancer)
- *Khc^mutA^* (Winding et al., 2016; Vladimir Gelfand; confirmed by lethality of hetero-allelic *Khc^mutA/8^* animals)
- *Khc^E177K^* and *Khc^E177K,R947E^* (Kelliher et al., 2018; Jill Wildonger)
- *Df(Khc) (Df(2R)BSC309*; Cook et al., 2012; BL #23692)
- *milt^92^* (Cox and Spradling, 2006; Stowers et al., 2002; Tom Schwarz)
- *Df(milt) (Df(2L)ED440, P{w[+mW.Scer\FRT.hs3]=3’.RS5+3.3’}ED440*; Ryder et al., 2004; Kyoto #150498)
- *Miro^Sd32^* (Guo et al., 2005)
- *MiroB^682^* (Guo et al., 2005; BL #52003)
- *Df(Miro) (Df(3R)Exel6197*; Parks et al., 2004; BL #7676)
- *Klc^8ex94^* (Gindhart et al., 1998; BL #31997)
- *syd^z4^* (Bowman et al., 2000; BL #32016)
- *Df(3L)sydA^2^* (Bowman et al., 2000; deleting C-terminus; BL #32017)
- *Pat1^gnve^* and *Pat1^robin^* (Loiseau et al., 2010; Isabel Palacios)
- *Sod1^n1^* (Phillips et al., 1989; BL #24492)
- *Sod1^n64^* (Phillips et al., 1995; BL #7451)
- *Rtnl1-YFP (PBac{681.P.FSVS-1}Rtnl1CPTI001291*; Cahir O’Kane; O’Sullivan et al., 2012)
- *Drp1^T26^* (Verstreken et al., 2005; BL #3662)
- *MarfB* (Sandoval et al., 2014; Hugo Bellen)
- *P{lacW}Opa1^s3475^* (Spradling et al., 1999; BL #12188)
- *Cat”^1^* (Mackay and Bewley, 1989; Matthias Landgraf)
- *Sod2^n283^* (Duttaroy et al., 2003; BL#34060)
- *Pex3^2^* (Faust et al., 2014; BL#64251)

Gal4 driver lines used were the

- *elav-Gal4* (Luo et al., 1994)
- tubP-Gal4 (Lee and Luo, 1999; Liqun Luo)

UAS lines

- *UAS-Khc^FL^::GFP* (3^rd^, unpublished; Isabel Palacios)
- *UAS-Khc^82-711^-GFP* (2^nd^, BL #9648; constitutively active Khc consisting of base pairs 248-2134 / aa 82-711; fused with EGFP sequence; flybase.org: FBrf0198610)
- *UAS-Khc1-850-GFP* (Loiseau et al., 2010; Isabel Palacios)
- *UAS-Khc-RNAi* (Lu et al., 2013; Vagnoni et al., 2016; BL #35770)
- *UAS-Sod1* (J. Hu and J.P. Phillips, unpublished)
- *UAS-Duox* (Ha et al., 2005; Matthias Landgraf)
- *UAS-Gapdh-IR* (*Gapdh1^GD7467^*; Vienna *Drosophila* Resource Centre)

### Cloning of *UAS-Nox-YPet*

*10xUAS-IVS-Nox::YPet* was generated by using the *pJFRC12-10XUAS-IVS-myr-GFP* vector (Addgene 26222; Pfeiffer et al., 2010) as a backbone which was modified by substituting GFP with YPet (Nguyen and Daugherty, 2005) plus an N-terminal flexible linker, amplified from dFlex_YPet_phase0 (Gärtig et al., 2019) using primer ML1 and ML2, and inserted by the Klenow Assembly Method (tinyurl.com/4r99uv8m) into the XbaI/BamHI sites producing Vector 1: *pJFRC12-10xUAS-IVS-myr-linker-YPet*. Nox cDNA was amplified from a DGRC (*Drosophila* Genomics Resource Center) cDNA library clone using primers ML5 and ML6, located in a *pOTB7* vector backbone, and inserted into BamHI/XhoI sites of Vector 1. Constructs were sent to FlyORF for transgenesis, and targeted via PhiC31-mediated site-specific insertion to the *PBac{y^+^-attP-3B}VK00040* landing site (Bloomington line #9755) on the third chromosome (3R, 87B10).

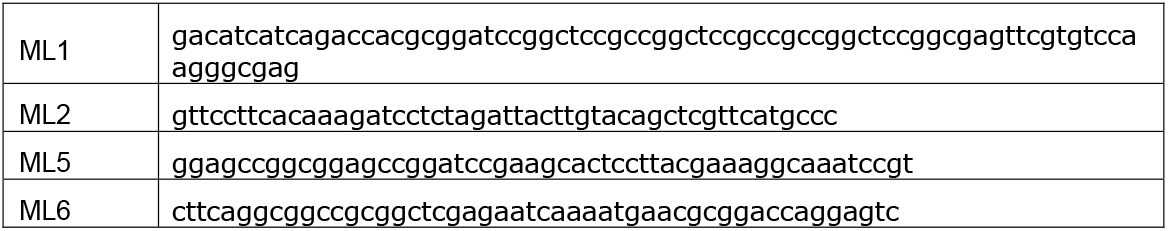

### *Drosophila* primary cell culture

*Drosophila* primary neuron cultures were performed as published previously (Prokop et al., 2012; Qu et al., 2017). In brief, stage 11 embryos were treated for 1 min with bleach to remove the chorion, sterilized for ~30 s in 70% ethanol, washed in sterile Schneider’s/FCS, and eventually homogenized with micro-pestles in 1.5 centrifuge tubes containing 21 embryos per 100μl dispersion medium and left to incubate for 5 min at 37°C. Cells were washed with Schneider’s medium (Gibco), spun down for 4 mins at 650g, supernatant was removed and cells re-suspended in 90μl of Schneider’s medium containing 20% fetal calf serum (Gibco). 30μl drops were placed on cover slips. Cells were allowed to adhere for ~2hrs either directly on glass or on cover slips coated with a 5 μg/ml solution of concanavalin A, and then grown as a hanging drop culture for hours or days at 26°C as indicated in each experiment.

To abolish maternal rescue of mutants, i.e. masking of the mutant phenotype caused by deposition of normal gene product from the healthy gene copy of the heterozygous mothers in the oocyte (Prokop, 2013), we used a pre-culture strategy (Prokop et al., 2012; Sánchez-Soriano et al., 2010) where cells were kept for 5 days in a tube before they were plated on a coverslip.

Cells were treated with 100 μM Trolox (Sigma; stepwise diluted from a 100mM stock solution in ethanol) or 100 μM DEM prepared in 100% ethanol. For controls (vehicle treatment), equivalent concentrations of vehicle (sterile H2O or 100% ethanol) were diluted in cell culture medium. All reagents were purchased from Sigma-Aldrich, unless otherwise stated.

For visualisation of mitochondria, cell cultures were incubated with 400nM MitoTracker Red CMXRos (Invitrogen; Klionsky et al., 2012) for 30min at room temperature (RT); stock solutions were prepared in DMSO and diluted in cell culture medium to the final concentration. Following incubation, cultures were then fixed and stained following the procedures below.

### Immunohistochemistry

Primary fly neurons were fixed in 4% paraformaldehyde (PFA) in 0.05M phosphate buffer (PB; pH 7–7.2) for 30min at room temperature (RT). Antibody staining and washes were performed with PBT. Staining reagents: anti-tubulin (clone DM1A, mouse, 1:1000, Sigma; alternatively, clone YL1/2, rat, 1:500, Millipore Bioscience Research Reagents); anti-Syt (1:1000; rabbit; Sean Sweeney); anti-GFP (1:500, rabbit, ab290, Abcam); Cy3-conjugated anti-HRP (goat, 1:100, Jackson ImmunoResearch); FITC-, Cy3- or Cy5-conjugated secondary antibodies (1:200; donkey, purified, Jackson Immuno Research); F-actin was stained with Phalloidin conjugated with TRITC/Alexa647, FITC or Atto647N (1:200; Invitrogen and Sigma). Specimens were embedded in ProLong Gold Antifade mounting medium.

### Microscopy and data analysis

Standard documentation was performed with AxioCam monochrome digital cameras (Carl Zeiss Ltd). mounted on BX50WI or BX51 Olympus compound fluorescent microscopes. To determine the degree of MT disorganisation in axons we used the “MT disorganisation index” (MDI) (Qu et al., 2017): the area of disorganisation was measured using the freehand selection tool in Fiji/ImageJ; this value was then divided by axon length (see above) multiplied by 0.5 μm (typical axon diameter, thus approximating the expected area of the axon if it were not disorganised). To quantify the number of synaptic densities in mature neurons in culture, we used ImageJ, first thresholding to select synaptic densities from axons of single isolated cells, followed by particle analysis. For statistical analyses, Kruskal–Wallis one-way ANOVA with *post hoc* Dunn’s test or Mann–Whitney Rank Sum Tests were used to compare groups. The data used for our analyses will be made available on request from the authors.

## Ethical statement

An ethical statement is not required.

---

**Fig.S1.**
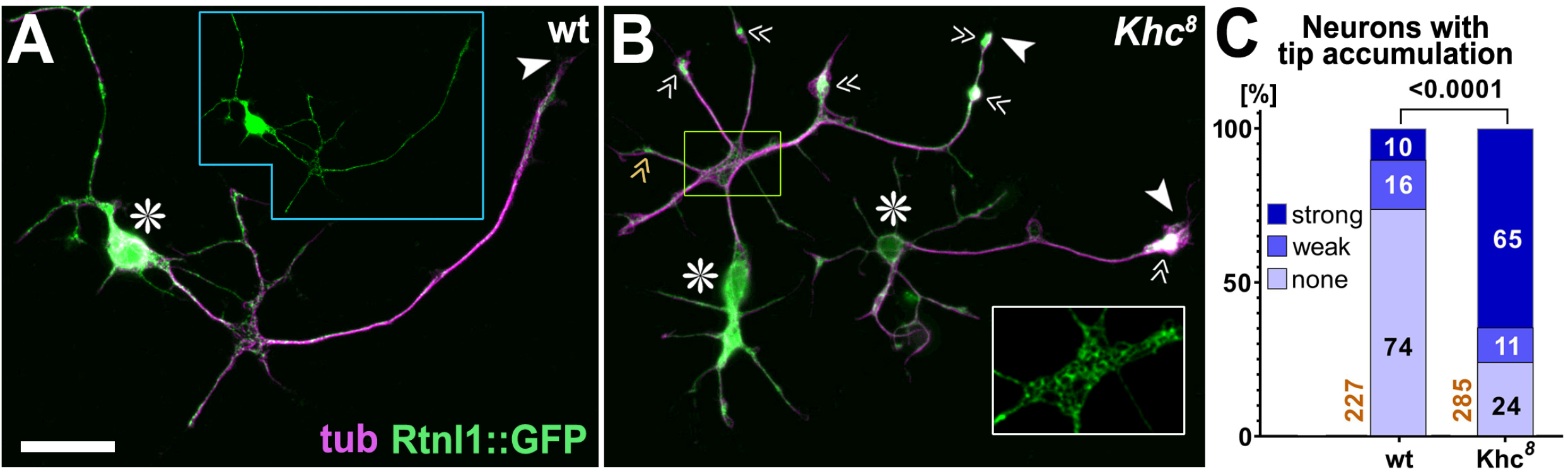
Endoplasmatic reticulum accumulates at axon tips upon loss of Khc. **A,B**) Primary neurons at 5 DIV carrying the genomically tagged *Rtln1-YFP* allele labelling endoplasmic reticulum (ER; del Castillo et al., 2019; O’Sullivan et al., 2012), either in wild-type (wt; A) or *Khc^8^* mutant background (B); inset with blue outline in A displays the green channel of the neuron (reduced to 50% in size) to illustrate the continuous nature of Rtnl1::GFP-labelled ER throughout its neurites; the yellow emboxed area in B is shown as twofold increased inset of the green channel to illustrate the netlike organisation of ER visible in axonal swellings. Asterisks indicate cell bodies and arrow heads axon tips (note that there are two neurons in B), white/orange chevrons point at strong/weak axonal tip accumulations of ER. Accumulations might indicate an imbalance of antero- and retrograde organelle movement potentially caused by loss of Khc-dependent Dynein transport to axon tips (Moughamian et al., 2013; Twelvetrees et al., 2016) expected to reduce the retrograde drift of ER. The scale bar in A represents 20 μm in A and B. **C**) Quantification of axonal tip accumulation of ER: numbers of neurons analysed are shown in orange, numbers in bars the rounded percentages of neurons with no/weak/strong accumulations, the number above bars show the P value of the X^2^ test.

**Fig.S2.**
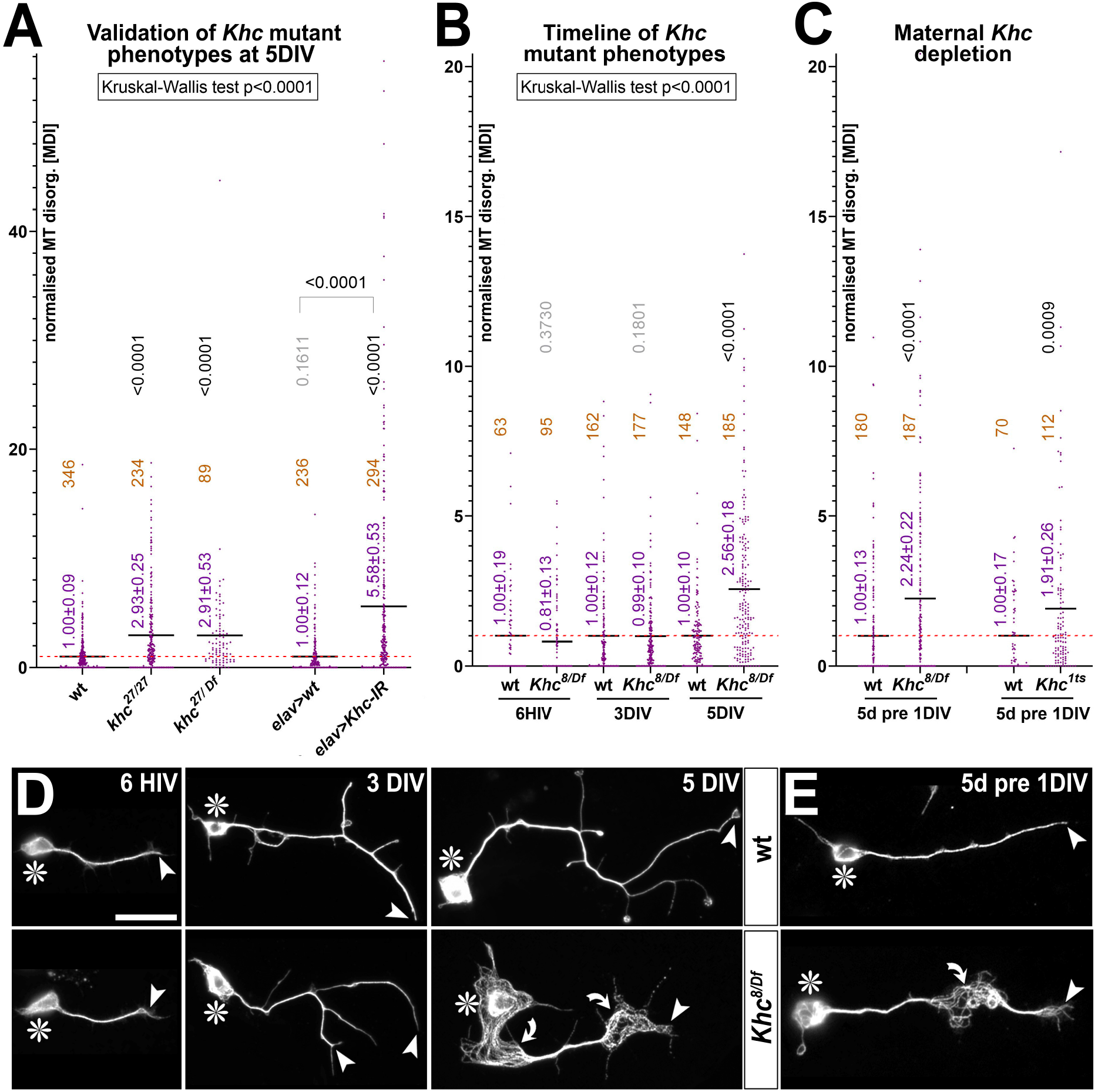
Validation of Khc’s MT phenotype and demonstration of maternal contribution. **A-C**) Quantification of MT curling phenotypes measured as MT disorganisation index (MDI) and normalised to wild-type controls (red stippled line); mean ± SEM is indicated in blue, numbers of analysed neurons in orange, results of Mann Whitney rank sum tests are shown in grey/black; A) shows data for *Khc^27^* in homozygosis (*27/27*) or over deficiency (*27/Df*), for Khc knock-down (*elav>Khc-IR)* and wild-type (wt) and driver line (*elav*) controls; B) shows data for *Khc^8^* over deficiency (*8/Df*) and wild-type controls at different culture times (HIV, hours *in vitro*; DIV, days *in vitro*); C) shows data for *Klc^8/Df^* and *Klc^1ts^* at 1DIV following 5d pre-culture; note that *Klc^1ts^* is a temperature-sensitive allele (see methods) and was pre-cultured at 26°C and cultured at 29°C. **D,E**) Examples of neurons at different times in culture (D; relating to data in B) and after pre-culture (E; relating to C); asterisks indicate cell bodies, arrow heads axon tips, curved arrows areas of MT curling; scale bar in D represents 20μm in D and E.

**Fig.S4.**
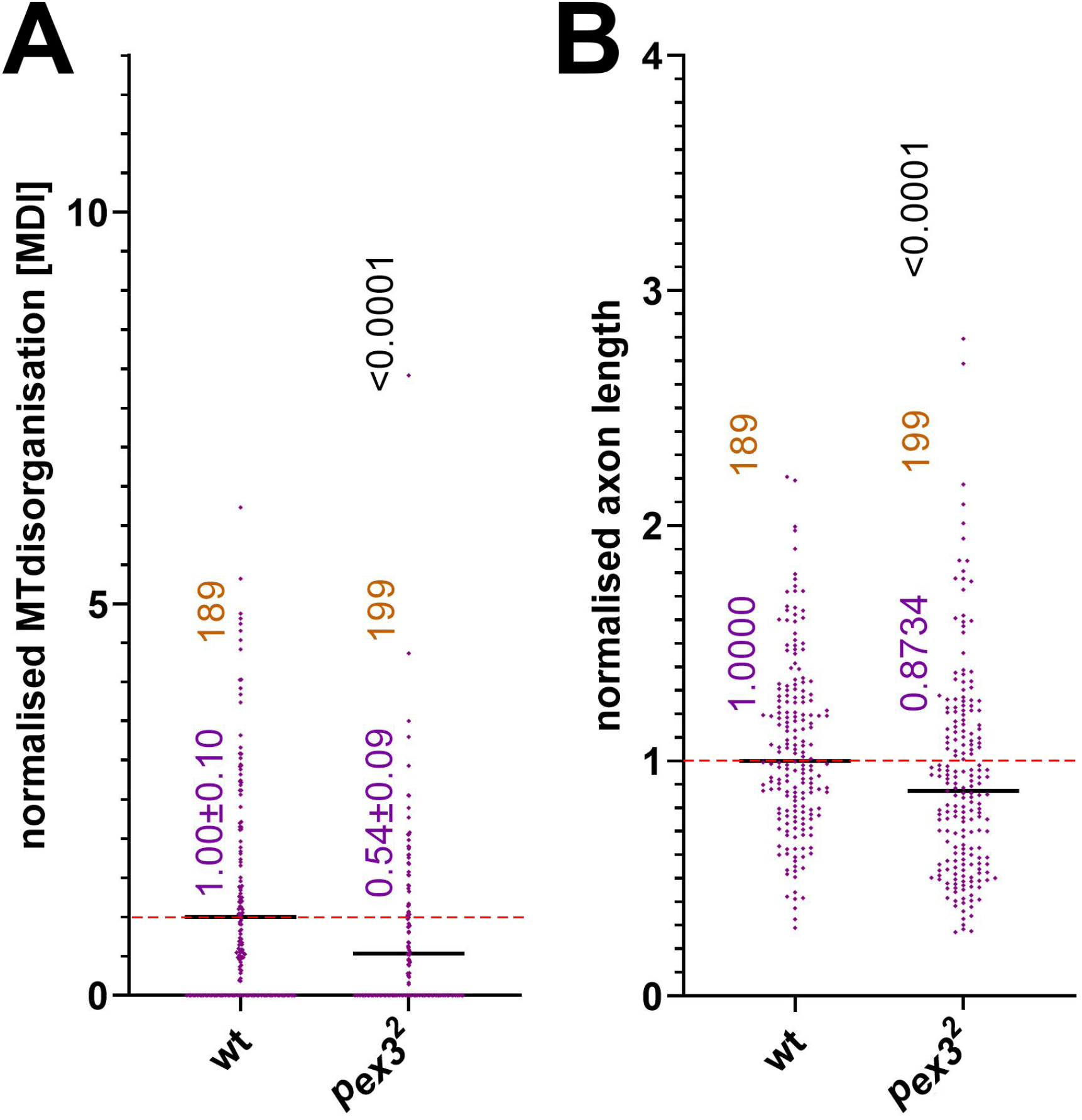
MT bundle and axon length phenotypes of *pex3^2^* mutant neurons. Quantification of phenotypes of wild-type (wt) and *Pex^3^* homozygous mutant neurons: **A**) MT curling phenotypes measured as MT disorganisation index (MDI); **B**) axon length; both measured are normalised to wild-type controls (red stippled line); mean ± SEM is indicated in blue, numbers of analysed neurons in orange, results of Mann Whitney rank sum tests are shown in grey/black.

**Fig.S5.**
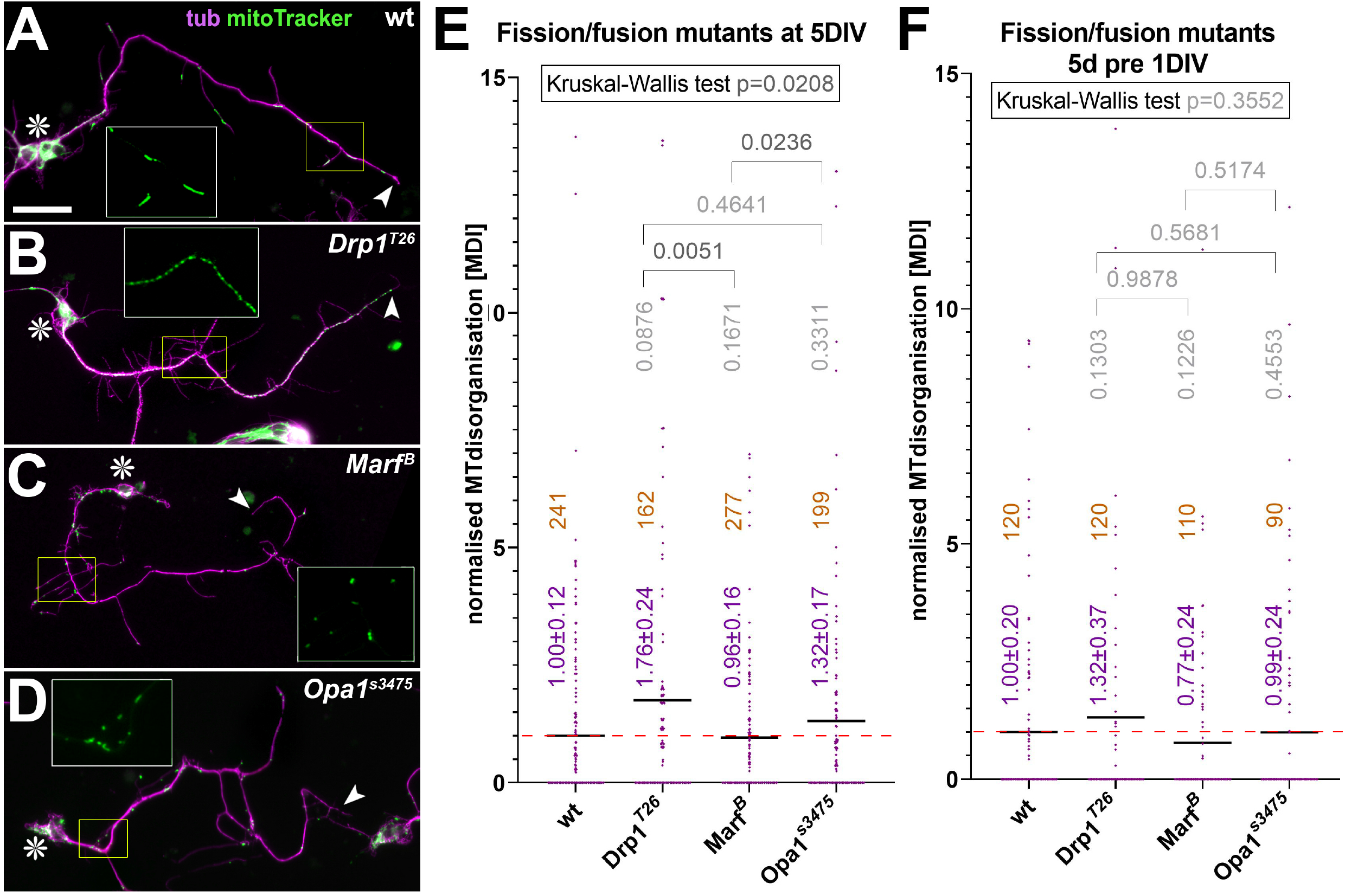
Impaired fission/fusion of mitochondria does not affect MT bundles. **A-D**) Neurons at 5 days *in vitro* (DIV) and stained with anti-tubulin (tub, magenta) and mitoTracker (green); they are wild-type (wt) or homozygous mutant for the mitochondrial fission factor Drp or the mitochondrial fusion factors Marf or Opa, as indicated; asterisks indicate cell bodies, arrow heads axon tips, yellow emboxed areas are shown as 2-fold enlarged insets (green channel only), and the scale bar in A represents 20μm in A-D; note that mitochondria tend to appear as dashed lines in controls (A), as a continuous string of pearls excluded from side branches upon loss of fission (B), and as sparse dots upon loss of fusion (C,D). **E**) Quantification of MT curling phenotypes from experiments shown in A-D, measured as MT disorganisation index (MDI) and normalised to wild-type controls (red stippled line). **F**) Similar experiments with the same mutations using 5 day pre-culture and culture for 1 day. In E and F, mean ± SEM is indicated in blue, numbers of analysed neurons in orange, results of Mann Whitney rank sum tests are shown in grey/black.

**Fig.S3.**
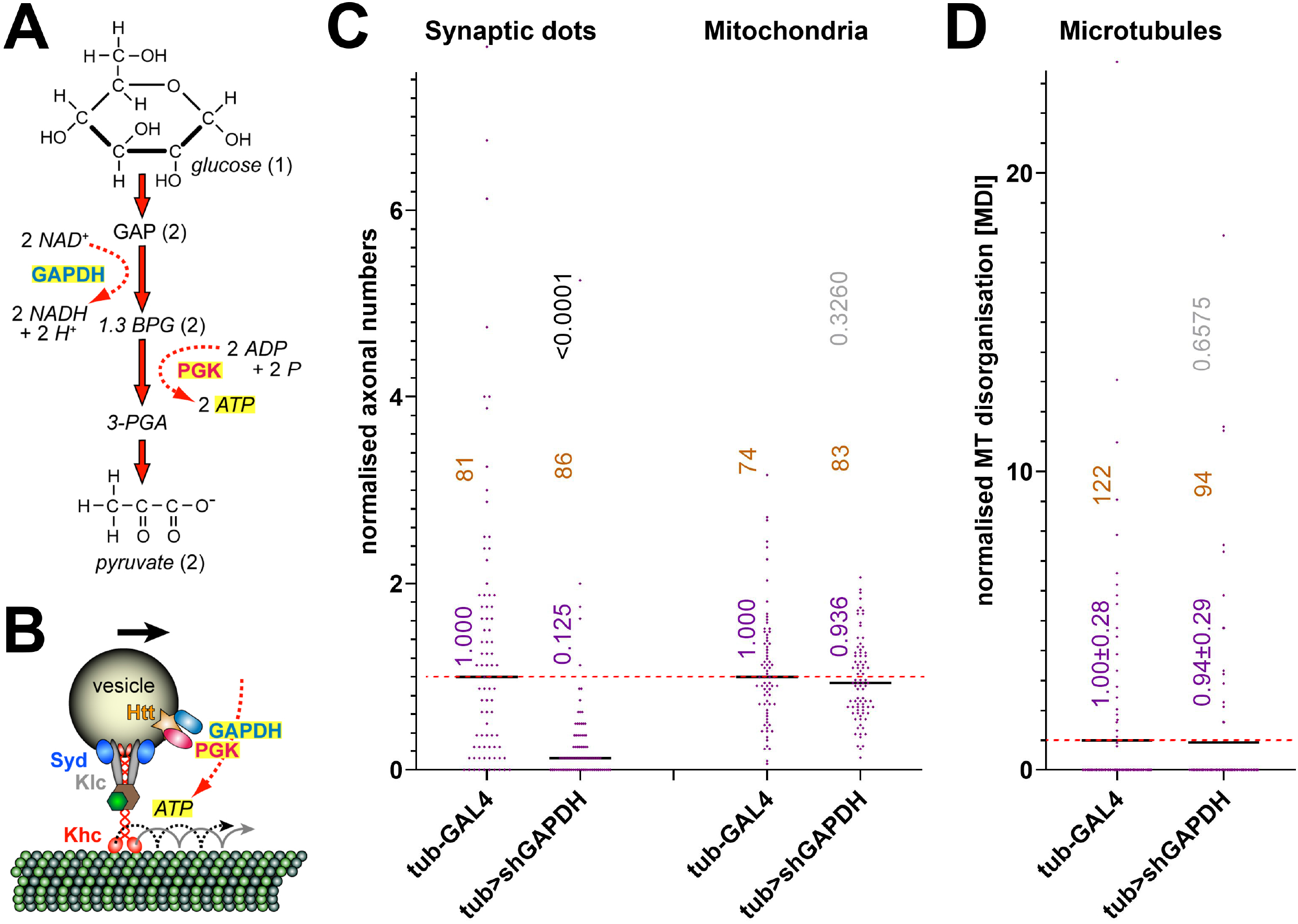
Phenotypes upon *Gapdh1* knock-down in primary neurons at 5 DIV. **A**) Illustration of the NADH- and ATP-generating steps of glycolysis; names of proteins are shown in bold, other molecules in italics: GAPDH (glyceraldehyde-3-phosphate dehydrogenase), PGK (phosphoglycerate kinase), GAP (glyceraldehyde-3-phosphate), 1,3 BPG (1,3-biphosphoglycerate), 3-PGA (3-phosphoglycerate). **B**) GAPDH and PGK are present on transported vesicles together with other factors relevant for glycolysis (Hinckelmann et al., 2016; Zala et al., 2013) providing ATP to drive kinesin-mediated processive transport (red and stippled black lines). **C**) In the absence of Gapdh1, the transport of synaptic vesicles but not mitochondria is impaired (assessed via anti-Syt and mitoTracker staining; see Fig.2), as is consistent with *in vivo* observations in *Drosophila* (larval motor nerves; Zala et al., 2013). **D**) Absence of GAPDH does not cause MT curling. Quantification of MT curling phenotypes in C and D is measured as MT disorganisation index (MDI) and normalised to wild-type controls (red stippled line); median in C and mean ± SEM in D are indicated in blue, numbers of analysed neurons in orange, results of Mann Whitney rank sum tests are shown in grey/black.

**Fig.S6.**
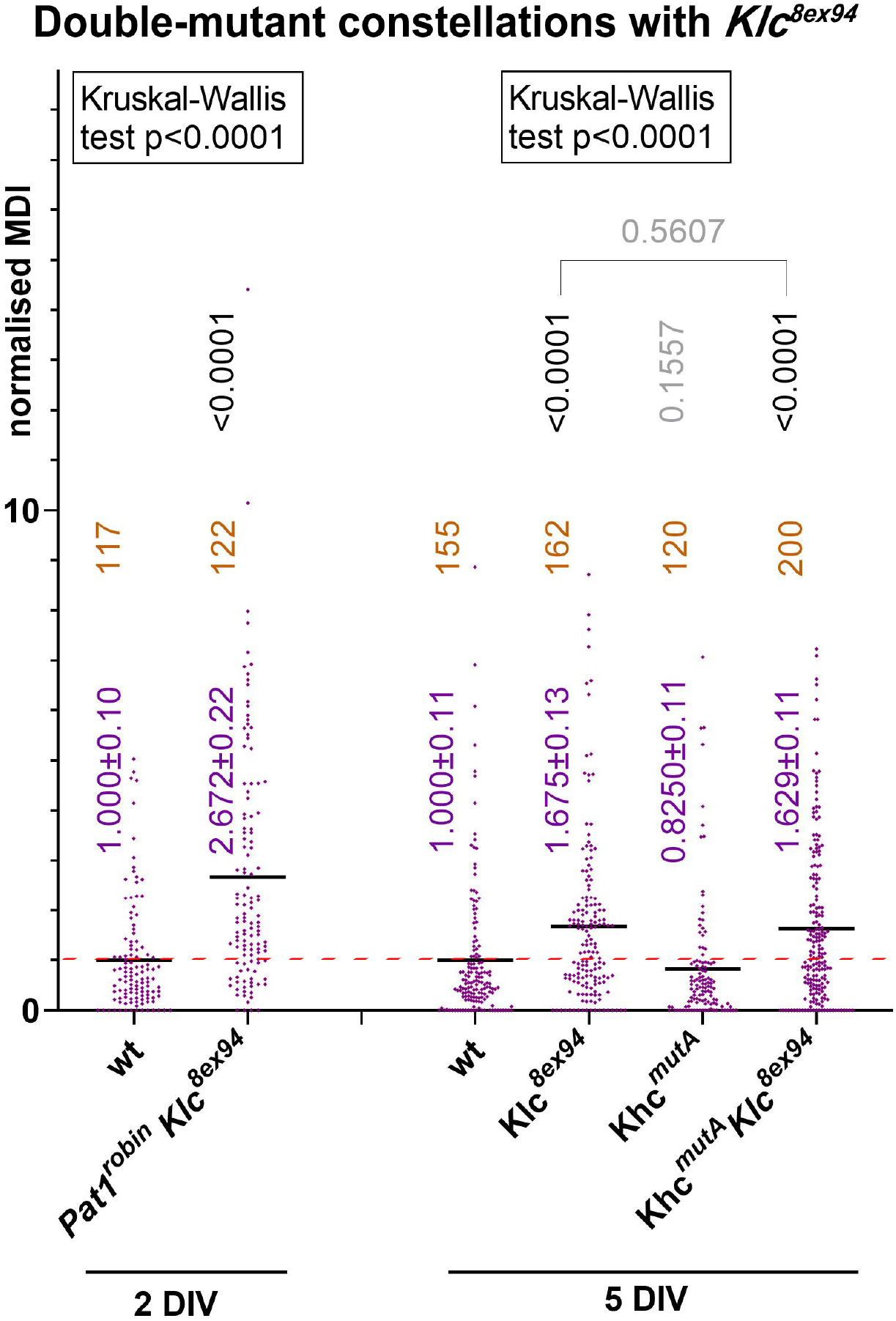
*Pat1* and *Khc^mutA^* mutations fail to suppress the Klc-deficient MT curling phenotype. Quantification of MT curling phenotypes measured as MT disorganisation index (MDI) and normalised to wild-type controls (red stippled line); genotypes are shown below, also indicating the culture period (2DIV, 5DIV); mean ± SEM is indicated in blue, numbers of analysed neurons in orange, results of Mann Whitney rank sum tests are shown in grey/black. The fact that *Khc^mutA^* and *Pat1^robin^* fail to suppress *Klc^8ex94^*-induced MT curling suggests that potential surplus pools of non-inactivated Khc do not engage in MT sliding or Pat1-mediated transport to cause MT curling.

## Notes

### Competing Interest Statement

The authors have declared no competing interest.

